# NAC transcription factor RD26 is a regulator of root hair morphogenic plasticity

**DOI:** 10.1101/2021.04.21.440803

**Authors:** Iman Kamranfar, Salma Balazadeh, Bernd Mueller-Roeber

## Abstract

Root hairs are outgrowths of epidermal cells central for water and nutrient acquisition. Root hair growth is plastically modified by environmental cues. A frequent response to water limitation is active shortening of root hairs, involving largely unknown molecular mechanisms. A root hair-specific *cis*-regulatory element (RHE) integrates developmental cues with downstream signalling of root hair morphogenesis. Here, we demonstrate NAC transcription factor RD26 to be a key expressional regulator of this drought stress-triggered developmental response in *Arabidopsis thaliana*. RD26 directly represses *RSL4* and *RSL1*, two master transcription regulators of root hair morphogenesis, by binding RHE. RD26 further represses core cell wall modification genes including expansins (*EXPA7*, *EXPA18*), hydroxyproline-rich glycoproteins (*LRX1*), xyloglucan endotransglucosylases/hydrolases (*XTH12*, *13*, *14*, *26*), class III peroxidases (*PRX44*) and plasma membrane H^+^-ATPase (*AHA7*) through RHE. Of note, several RD26-repressed genes are activated by RSL4. Thus, by repressing RSL4 and numerous cell wall-related genes, RD26 governs a robust gene regulatory network for restricting root hair growth under drought. A similar regulatory network exists in tomato, indicating evolutionary conservation across species.

**Significance statement:** In plants, root hairs play a vital role for water and nutrient acquisition, soil anchorage, and microbial interactions. During drought stress, root hair growth is suppressed as an adaptive strategy to save cellular energy. We identified NAC transcription factor RD26 as a key regulator of this developmental plasticity in the model plant *Arabidopsis thaliana*. RD26 directly and negatively controls the transcriptional activity of key root hair developmental genes, *RSL1* and *RSL4*. Furthermore, RD26 suppresses the expression of several functional genes underlying root hair development including numerous cell wall-related genes. RD26 thus governs a robust gene regulatory network underlying the developmental response to drought stress. A similar regulatory network exists in tomato indicating evolutionary conservation of this mechanism across species.

## Introduction

Plant roots exhibit a high degree of phenotypic plasticity which affects branching patterns (i.e., the number and positioning of lateral roots), and root hair density and length (1, 2). As plants are sessile, plasticity of root system architecture (RSA) is an important adaptive trait allowing plants to survive under different environmental conditions. The *Arabidopsis thaliana* root epidermis is composed of a single layer of cells organized in rows (files) of root hair cells or non-hair cells (3, 4). The fate-determining developmental signals are executed by different core transcription factors (TFs) which control the expression of downstream genes involved in the morphogenesis of root epidermal cells. The expression of the core fate-determining TFs is not much affected by environmental cues (5, 6).

The TF GLABRA2 (GL2) conveys signals from cell fate determination to cell differentiation (7). One of the key downstream genes of GL2 is *ROOT HAIR DEFECTIVE6* (*RHD6*) which encodes a bHLH TF involved in the regulation of root hair outgrowth. Three phylogenetically related bHLH TFs, *i.e.* ROOT HAIR DEFECTIVE 6-LIKE 1 (RSL1), RSL2 and RSL4, also promote root hair growth; *RSL4* and *RSL2* act downstream of RHD6 and RSL1 (8–11). RSL4 induces several genes involved in cell signaling, vesicle transport and cell wall modification (9, 12, 13). GL2 suppresses root hair development in non-hair cells by direct repression of *RHD6*, *RSL1* and *RSL2* (7).

The polar tip growth of root hairs requires a directed delivery of cell wall and membrane material to the growing tip, accompanied by cell wall loosening and reassembly. Expansins (14) and xyloglucan endotransglycosylase activities (15) are responsible for major cell wall modifications in growing root hairs.

Root hairs greatly increase the absorptive surface of the root, facilitating the uptake of nutrients. Phosphate deprivation stimulates root hair formation (16), while drought represses root hair growth (17). Similarly, salt stress decreases hair cell density and root hair length in a dose-dependent manner, leading to reduced ion uptake (18). Although the signalling networks controlling root hair development under non-stress conditions and in response to phosphate deprivation are well characterized, the gene regulatory networks repressing root hair growth during drought and salt stress are not yet well understood.

Transcription factors of the NAC (NAM, ATAF and CUC) family regulate diverse developmental processes and stress responses (19, 20), and they played an important role in land plant evolution (21). RD26 (ANAC072) is a NAC TF involved in stress signalling in *Arabidopsis thaliana*; together with six other NACs, namely ANAC002/019/032/055/081 and 102, it forms the ATAF clade. Overexpression of *RD26* improves drought tolerance by enhancing and repressing, respectively, ABA and BR signalling (22–24).

Here, we demonstrate that RD26 negatively regulates root hair growth downstream of ABA signalling by directly repressing *RSL1* and *RSL4* expression. RD26 also represses root hair outgrowth by direct repression of several root hair-specific genes involved in cell wall modifications. RD26 thus regulates developmental plasticity of the root in response to environmental cues. A similar regulatory network appears to be present in tomato, suggesting evolutionary conservation of the abiotic stress-responsive root hair developmental pathway across species.

## Results

### Expression of *RD26* is strongly affected by abiotic stresses in roots

The important role of roots for responses to abiotic stresses prompted us to investigate the expression of *ATAF* clade *NAC* genes in roots, utilizing public transcriptome data. *ATAF*s are strongly induced in roots by salinity and osmotic stress, while expression of the two phylogenetically most closely related *NACs*, *ANAC042* and *ANAC047*, was affected to a lesser extent (**Fig. 1a)**. The induction by multiple abiotic stresses in roots is particularly evident for *RD26*, which exhibited highest induction in salt, osmotic, heat, drought and cold stresses. Importantly, *RD26* appeared to be the only member of the *ATAF* family repressed by auxin treatment as well as by iron (Fe), sulphur and phosphate deficiency and acidification of the growth medium (**Fig. 1a**) which are all conditions leading to elongated root hairs. We also checked the expression of *RD26* in roots dissected into different organ parts after diverse treatments (25–27). *RD26* most strongly responded to the stresses in the root maturation zone: while salt stress highly induces *RD26* in this region, low pH, sulphur deficiency and auxin treatment repress it (**Supplementary Fig. 1**). Similarly, we observed enhanced activity of the *RD26* promoter in roots under salinity stress and after ABA treatment in a dose-dependent manner, mainly in non-meristematic regions (**Fig. 1b**). Expression of *RD26* in roots is enhanced after ABA, osmotic stress and salt treatment compared to control conditions (**Fig. 1c**), while shifting Arabidopsis seedlings to an acidic pH medium, or to iron, sulphur or phosphate deficiency reduces *RD26* expression compared to the control (**Fig. 1c**). We checked the accumulation of RD26 protein in *pRD26:RD26-GFP*/*rd26-1* seedlings expressing the RD26-GFP coding sequence from the native *RD26* promoter (28) by Western blot analysis which revealed elevated accumulation of RD26-GFP protein after 12 h of ABA treatment in a dose-dependent manner in root samples (**Fig. 1d**), Upon ABA treatment, RD26-GFP signal was detected by confocal microscopy in all inner and outer root layers mainly of the maturation zone (**Fig. 1e; Supplementary Video 1a, b**). In the root epidermis, GFP signal was much stronger in non-hair (N) cells than in hair (H) cells (**Fig. 1f**), and signal intensity was negatively correlated with the length of the root hairs (Pearson correlation coefficient = −0.63; **Fig. 1g**). The fact that *RD26* expression is enhanced by environmental conditions that shorten root hairs, but reduced by conditions that lead to longer root hairs strongly suggest a negative regulatory role of this TF in root hair plasticity.

**Figure 1.**
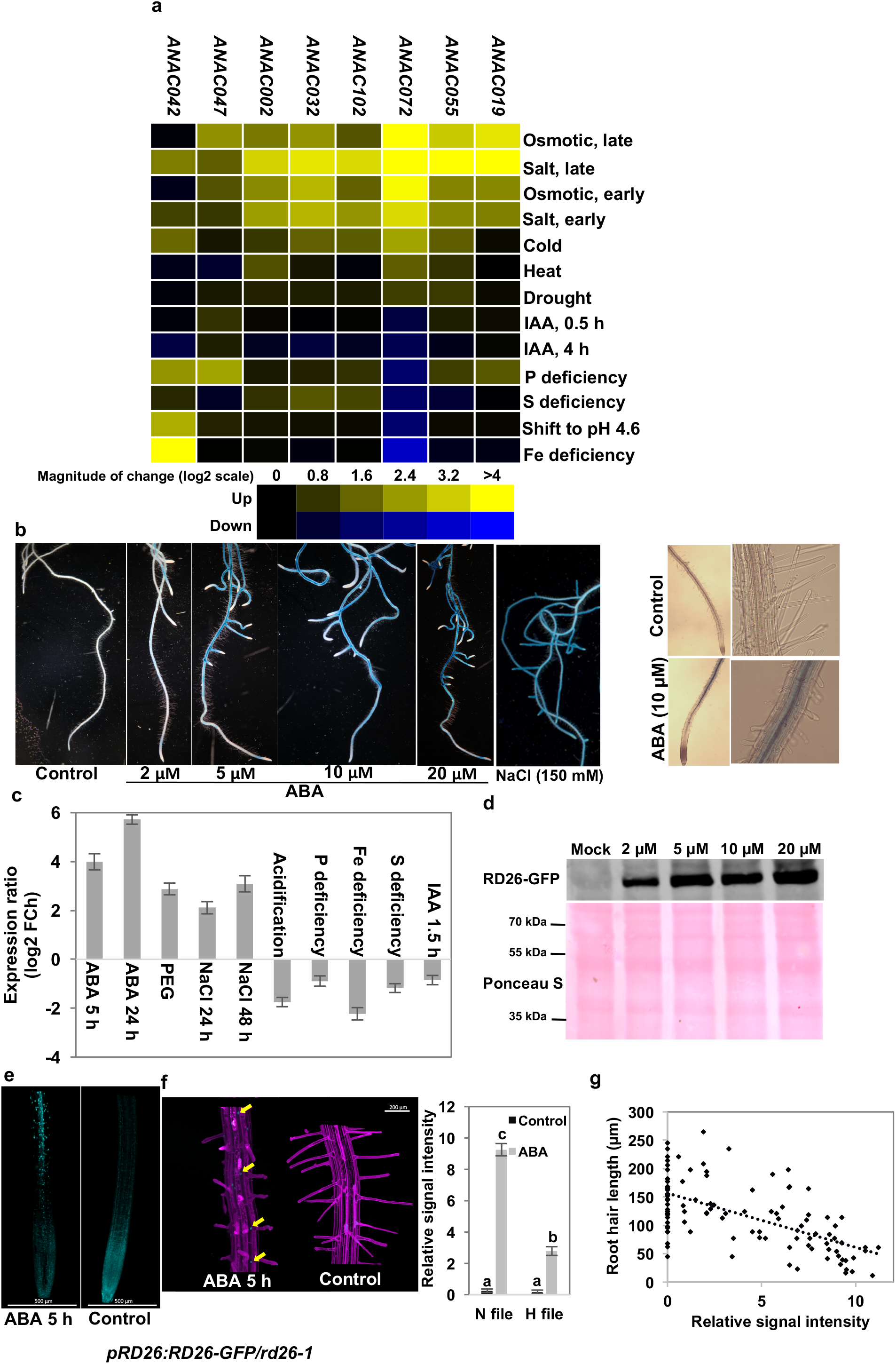
*RD26* is differentially expressed in roots under different abiotic stresses. **a,** Expression of ATAFs and the two closest ANACs in roots after different abiotic stresses or after auxin treatment. The heat map shows expression ratios of the genes in response to different treatments compared to control. Yellow, increased expression; blue, reduced expression. Data are from Genevestigator and represent the averages of at least three biological replicates each. **b,** Higher GUS activity in roots of *Pro_RD26_:GUS* plants in response to ABA and salt treatment. Two-week-old agar grown seedlings were transferred to liquid MS medium containing relevant chemicals and incubated for 3 h and then were stained. **c,** *RD26* transcript level determined by qRT-PCR in roots of agar-grown seedlings compared to control. For details of treatment see Methods. The data shown are means obtained from three biological replicates ± SD. **d,** Immunoblot analysis shows elevated levels of RD26-GFP protein in roots after 12 h ABA treatment compared to controls, in 10-day-old *pRD26:RD26-GFP*/*rd26* plants, using anti-GFP antibody (Western blot). kDa, kilo-Dalton. **e,** Confocal images show higher accumulation of RD26-GFP protein in nuclei of roots (mainly the maturation zone), and in **f,** files of non-hair cells (N file) than hair cells (H file) after 5 h ABA (15 µM) treatment. The relative signal intensity was quantified in 30 N and H cells per seedling; *n* = 20, one-way analysis of variance (ANOVA) least significant difference (LSD) test. Different letters indicate significant differences (*P* < 0.01). Roots were stained using propidium iodide to visualize walls of intact cells. **g,** Negative correlation between the length of root hairs and the intensity of the nuclear signal of RD26-GFP after 16 h ABA (15 µM) treatment. Relative signal intensity was quantified in five sequential cells in an H file in the maturation zone; *n* = 60. For f and g, *pRD26:RD26-GFP*/*rd26-1* six-day-old agar grown seedlings were transferred to liquid MS medium containing ABA, or ethanol as control.

### *RD26* suppresses growth of primary and lateral roots

ABA represses the growth of both, primary roots (PR) and lateral roots (LR) during stress (29). As ABA strongly enhances *RD26* expression in roots, we studied RSA in *RD26* transgenic lines (28). Root growth was reduced in *RD26* overexpressors (*RD26Ox*) compared to WT and an empty vector (*EV*) control line (**Supplementary Fig. 2, 3**). This decrease mainly resulted from a reduction of the number of emerged LRs (**Supplementary Fig. 2d, e**). *RD26Ox* lines were not significantly different than WT in the number of pre-emergent LRs, suggesting that RD26 mainly represses LR emergence rather than the formation of LR primordia (**Supplementary Fig. 2f, g; Supplementary Fig. 3a-c**). Investigating four transcriptome datasets^35-38^, we found that *RD26* is repressed during LR primordia development (**Supplementary Fig. 4**). The data support the model that RD26 acts as a negative regulator of LR development.

### *RD26* suppresses the growth of root hairs

Within the Brassicaceae and related families, plants facing drought stress adopt an ABA-dependent adaptive root growth strategy termed drought rhizogenesis whereby shortened, tuberized and hairless roots are formed (17, 30, 31). We confirmed the effect of ABA on drought rhizogenesis by growing Arabidopsis seedlings in the absence or presence of ABA (15 µM). ABA caused a 76% reduction in PR length compared to seedlings grown without ABA at 10 days after sowing (DAS; **Supplementary Fig. 5a, b**). Furthermore, ABA reduced both, the number (60%) and length (82%) of root hairs compared to control and it altered the shape of the root hairs from elongated to bulbous (**Supplementary Fig. 5a, c, d**).

The negative correlation between RD26-GFP protein level and root hair length (**Fig. 1g**), and the fact that *RD26* expression is strongly enhanced by ABA in roots, led us to assess the potential role of *RD26* for root hair formation, using transgenic plants expressing *RD26* from a β-estradiol (EST)-inducible promoter (*RD26-IOE* lines) (28). Seedlings transformed with the empty pER8 vector (*pER8-EV*) served as controls. Seedlings were grown for 5 days on ½ MS medium in the absence of ABA and EST, and then transferred to agar plates containing either ABA (10 µM) or EST (15 µM), or both (omitted in mock-treated seedlings), and root and root hair phenotypes were analysed 5 days after the transfer (DAT). ABA strongly elevated *RD26* expression in both lines (by ∼10-fold), 24 h after the treatment, while EST increased *RD26* transcript abundance only in *RD26-IOE* seedlings (by ∼6-fold), as expected. A combined treatment with both, ABA and EST enhanced *RD26* transcript abundance by ∼23-fold compared to mock treatment (**Fig. 2a**). ABA inhibited growth of PR in both lines, while EST inhibited PR growth only in *RD26-IOE* seedlings (by 61%) compared to mock-treated seedlings (**Fig. 2b, c**). A simultaneous treatment of *RD26-IOE* seedlings with EST and ABA further reduced PR length to 85% of that in mock-treated seedlings (**Fig. 2b, c**). Next, we analysed the number, length and density of root hairs. EST treatment reduced the number of root hairs by 61%, their length by 60%, and the density by 47% in *RD26-IOE* but not *pER8-EV* seedlings (**Fig. 2d-g**), while ABA treatment reduced all three traits in both lines. Simultaneous treatment with EST and ABA even more severely reduced root hair traits in *RD26-IOE* seedlings compared to mock-treated controls (**Fig. 2d, e**). Similarly, PR length and root hair number were reduced by 76% and 58% respectively, in *35S:RD26-GFP* seedlings compared to WT at 8 DAS (**Supplementary Fig. 3**).

**Figure 2.**
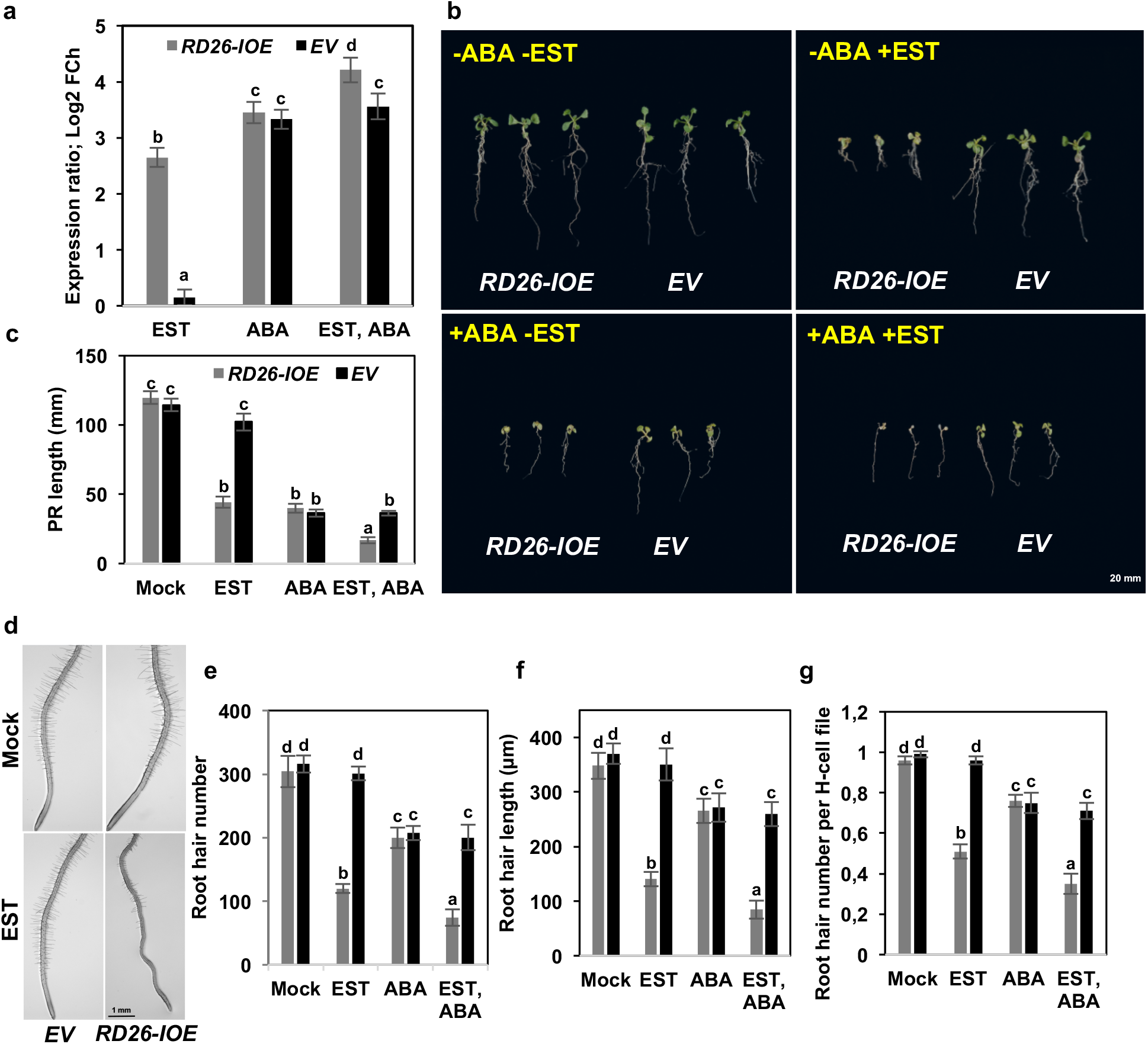
Induction of *RD26* mimics the ABA diminishing effect on the root and root hair. **a,** Expression of *RD26* in *RD26-IOE* and *EV* control plants after treatment with EST, ABA, or EST + ABA after 24 h incubation, as determined by qRT-PCR. **b,** Induction of *RD26* with EST, as well as treatment with ABA, represses root growth in *RD26-IOE* plants. **c,** ABA represses PR growth in *EV* and *RD26-IOE* plants. Induction of *RD26* expression by EST in *RD26-IOE* seedlings has a similar effect. **d - g,** ABA represses root hair formation in *EV* and *RD26-IOE* seedlings, while EST treatment inhibits the growth process only in *RD26-IOE* plants. Seedlings were transferred to new MS agar plats containing relevant chemicals at 5 DAS and analysed at 10 DAS (*n* = 20). Phenotyping of root hairs was performed for the first 10 mm to the root tip. Data are mean values obtained from three biological replicates ± SD. One-way analysis of variance (ANOVA) least significant difference (LSD) test. Different letters indicate significant differences (*P* < 0.01).

*ANAC019* and *ANAC055* are the closest homologs of *RD26* (32) indicating potential functional redundancy. We, therefore, tested the effect of ABA on root hair morphogenesis in *rd26* single and *rd26*/*anac019*/*anac055* triple knockout mutants (*triple nac* in the following). While in control condition knockout lines were not significantly different from WT with respect to root hair length and number, the inhibitory effect of ABA on root hair length in the WT was compromised in *rd26* and *triple nac* mutants (**Fig. 3a, b**). With respect to the number of root hairs, only *triple nac* mutant seedlings were less sensitive to ABA treatment than the WT (**Fig. 3c**). Furthermore, ABA-mediated repression of key genes involved in root hair outgrowth was less pronounced in *rd26* knockouts and particularly in *triple nac* seedlings (**Fig. 3d**).

**Figure 3.**
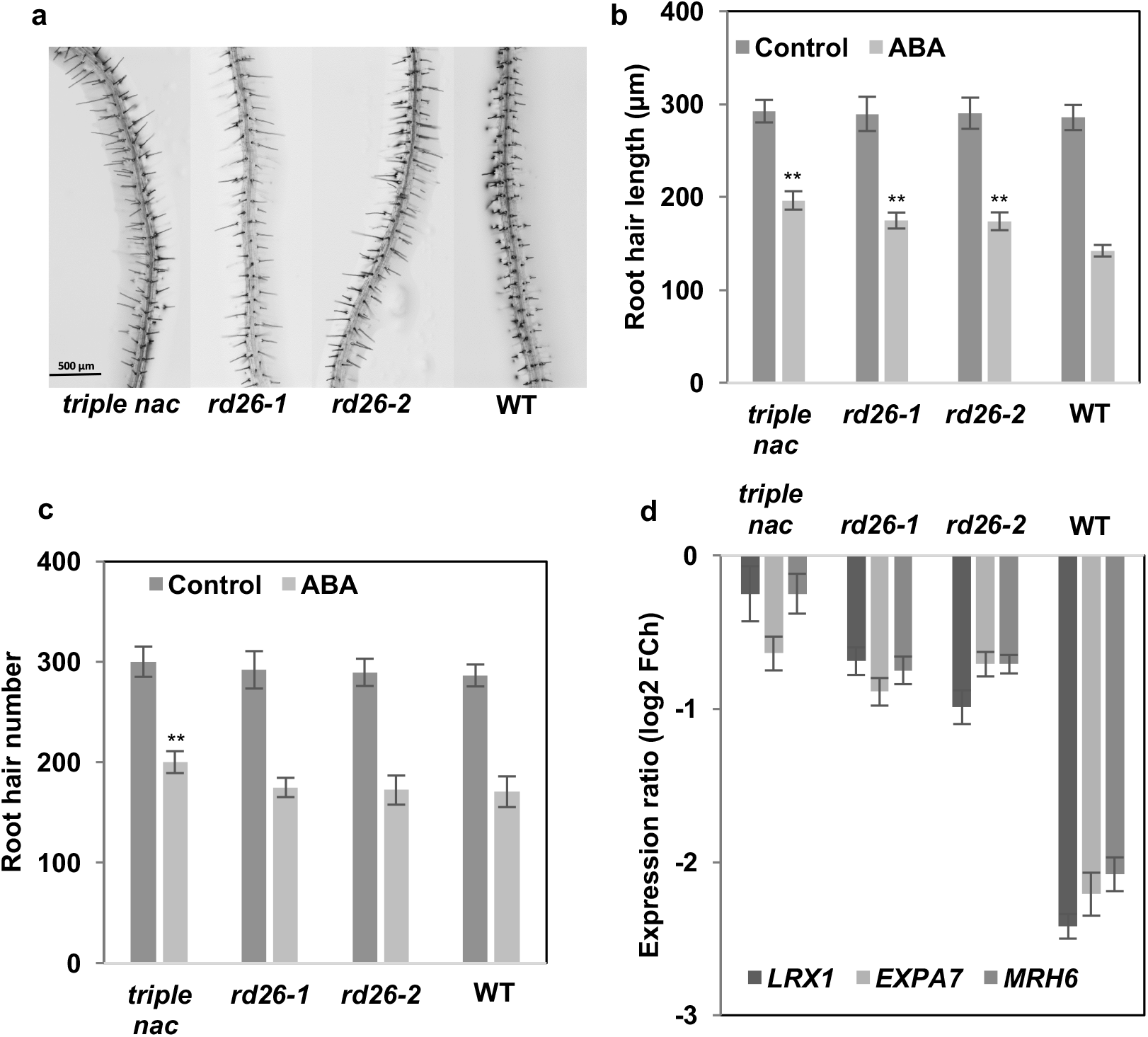
The inhibitory effect of ABA on root hair growth is compromised in *rd26* mutants. **a,** Agar-grown seedlings were transferred (at 5 DAS) to ABA- or mock-containing plates and photos to capture root hairs were taken three days later. **b,** Root hair length and **c,** root hair number at 8 DAS (*n* = 20). Root hairs were analysed within the first 10 mm to the root tip. **d,** Expression of root hair marker genes in WT seedlings and *rd26* and *triple nac* mutants. Date are given as the ratio of expression in ABA- *vs*. mock-treated seedlings. Note, that marker genes for root hair growth are less repressed by ABA treatment in mutants than in WT plants. Data are means of three biological replicates ± SD. Asterisks indicate a statistically significant difference from WT (ANOVA, ** *P* < 0.01).

### Effect of RD26 on root hair formation under osmotic stress conditions

Next, we tested root growth under conditions of osmotic stress imposed by treatment with polyethylene glycol (PEG) to decrease water potential. We transferred 4-day-old agar-grown seedlings to new plates infused with low (LC) or high concentration (HC) of PEG, or to PEG-free plates as control, and 6 days later checked the root hairs. In general, growth of root hairs was strongly inhibited in the presence of PEG (**Fig. 4a-c**). When WT seedlings were grown on LC plates, root hair number decreased by 55%, and root hair length decreased by 64% compared to seedlings grown on PEG-free medium. Importantly, in *rd26* knockout mutants the development of root hairs was significantly less affected by PEG treatment. On LC plates, root hair number in *rd26* mutants was reduced by only 37%, compared to growth on PEG-free plates. Similarly, root hair length was reduced by only 50% in both *rd26* knockout mutants compared to PEG-free conditions. When grown on HC plates, root hair number was reduced by 78% in WT, but by only 64% in *rd26* mutants.

**Figure 4.**
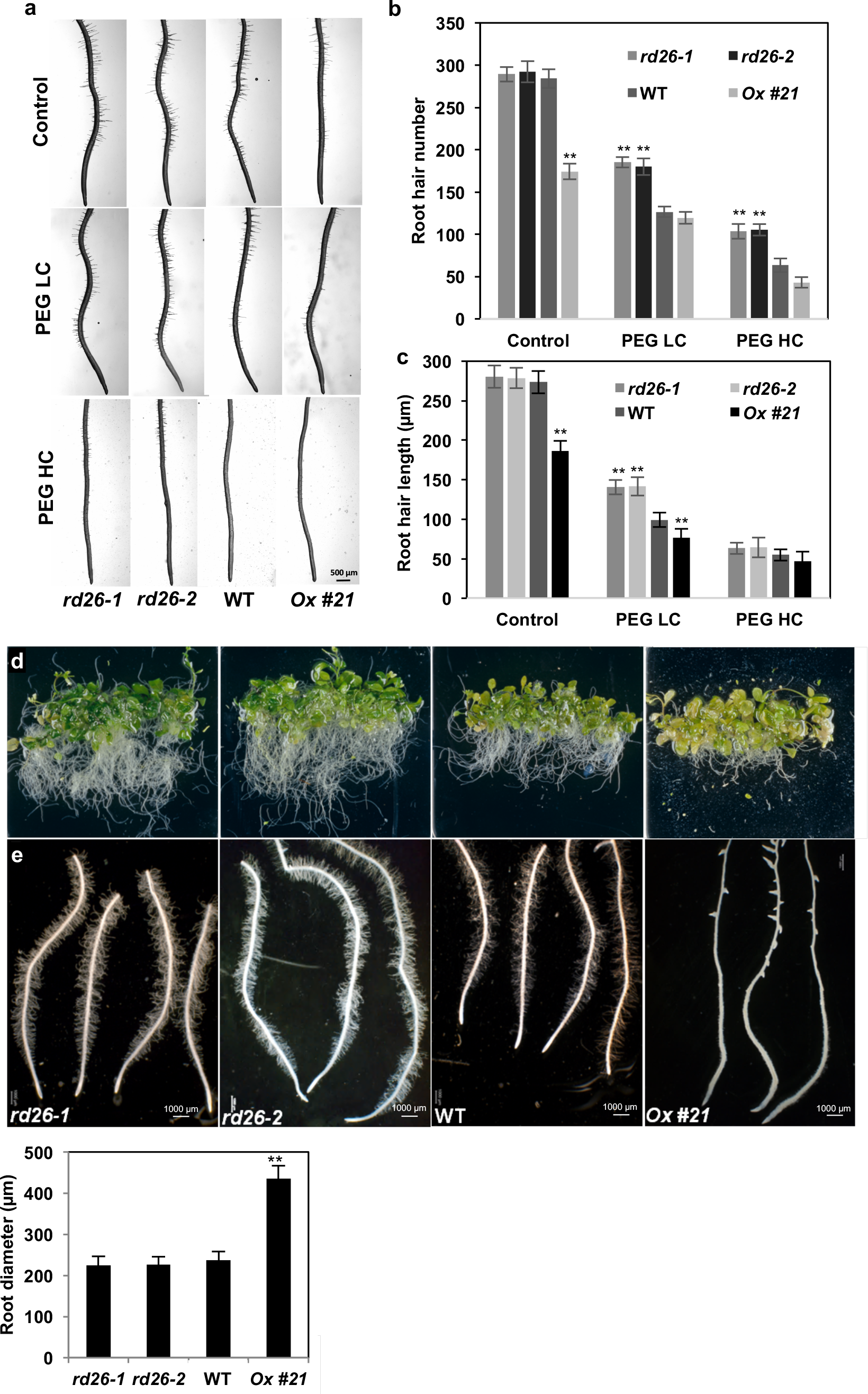
RD26 mediates drought rhizogenesis under osmotic stress conditions. **a,** Four-day-old agar-grown seedlings were transferred to new plates infused with PEG at low concentration (200 g/L PEG, LC plates, water potential −0.4 MPa) or high concentration (400 g/L PEG, HC plates, water potential −0.6 MPa), or to PEG-free plates as control (water potential −0.2 MPa), and 6 days later photos were captured and **b,** root hair number and **c,** length were studied in the first 10 mm to the root tip. **d,** Two-week-old agar-grown seedlings were transferred to liquid MS medium containing 250 g/L PEG (omitted in mock treatment) and incubated for 5 days. PEG repressed root growth more in *RD26ox* plants than in WT, while roots of *rd26* were largely insensitive to PEG treatment. **e,** Development of root hairs is negatively controlled by RD26 under PEG treatment and **f,** RD26 positively regulates root diameter under PEG treatment. Data are the means of two biological replicates ± SD (*n* > 20). Asterisks indicate statistically significant differences compared with WT (ANOVA, ** *P* < 0.01).

Next, two-week-old agar-grown Arabidopsis seedlings were transferred to liquid MS medium containing PEG and incubated for 5 days. PEG treatment repressed root growth in WT and more severely in *RD26Ox* seedlings, while the inhibitory effect of PEG was significantly reduced in *rd26* mutants (**Fig. 4d, e**). In *RD26Ox* seedlings, PEG treatment strongly inhibited root hair growth, repressed LR growth and significantly increased root diameter close towards the root tip (**Fig. 4f**). The observed growth phenotypes (i.e., repressed root growth, tuberized roots, and root hairlessness) are in accordance with drought rhizogenesis earlier reported (17). On the contrary, roots of *rd26* mutants produced numerous long root hairs, while WT roots exhibited an intermediate phenotype.

### RD26 is an RHE-binding factor

A bipartite, root hair-specific *cis*-element (RHE) is required for gene expression specifically in root hair cells in the maturation zone, and it was suggested that hormonal and environmental cues act downstream of transcription factor RHD6 to affect root hair growth *via* RHE (4, 5). The sequence motif ATCGTGNNNNGCACGTC is the most frequent version of RHE in root hair-expressed genes of angiosperms (5). We recently identified two related bipartite DNA binding sites (BS) of RD26, namely CGTr(n5-6)YACGyhayy (RD26-BSI) and rgwnDnY(n8-9)YACGtmwcy (RD26-BSII) (28). Both RD26 BSs well match the RHE *cis*-element (**Fig. 5a**), suggesting that RD26 might control its target genes as an RHE-binding factor.

**Figure 5.**
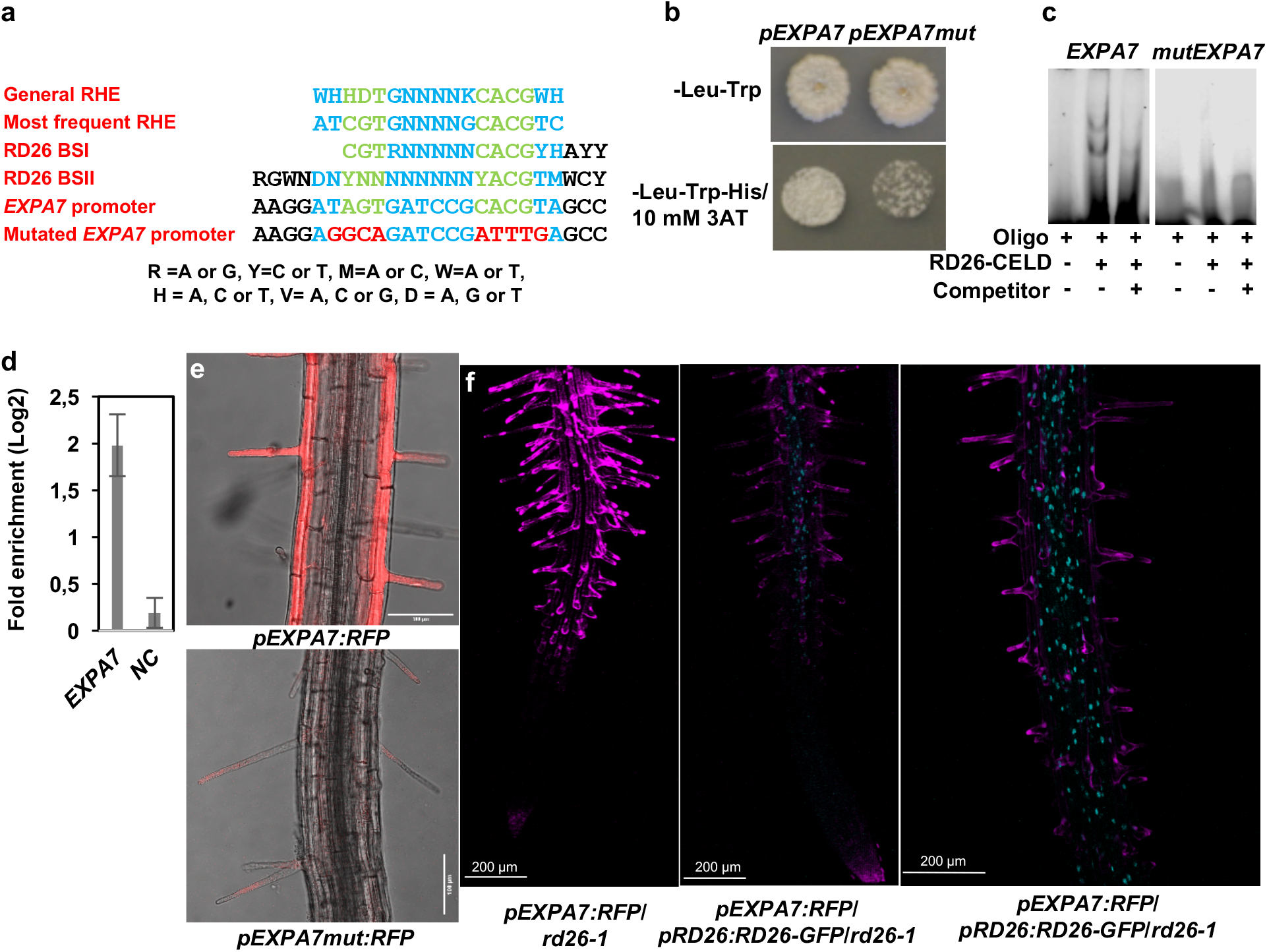
RD26 is an RHE-binding factor. **a,** The RD26 binding sites well match the RHE *cis*-element. **b,** Yeast one-hybrid assay confirmed physical interaction between RD26 and the promoter of *EXPA7*. The interaction diminished when a mutated version of the *EXPA7* promoter, lacking the RHE *cis*-element, was employed in the assay. **c,** EMSA. RD26-CELD protein binds specifically to the RD26 binding site (RHE) within the *EXPA7* promoter. DNA binding reactions were performed using RD26-CELD protein and a 40-bp promoter fragment containing the wild-type or a mutated RD26 binding site (sequences are given in panel a). Left panel: RD26-CELD binds to labelled wild-type double-stranded oligonucleotide (middle), while binding is not detected in the absence of RD26-CELD (left) or when non-labelled competitor is added (100-fold molar excess; right). Right panel: no binding to mutated *EXPA7* promoter is observed. **d,** ChIP-qPCR. *RD26:RD26-GFP*/*rd26-1* and *rd26-1* (as control) seedlings grown on ½ MS agar plates were transferred to ½ MS liquid culture and treated for 24 h with 15 µM ABA. The enrichment of the *EXPA7* promoter region containing the RD26 binding site was quantified using qPCR. As a negative control (*NC*), qPCR was performed on a promoter (*At2g22180*) lacking RD26 binding sites. Data represent means ± SD (two independent biological replicates, each with three technical replicates). **e,** RHE is required for the specific activity of the *EXPA7* promoter in root hair cells. The wild-type *EXPA7* promoter and a mutated version lacking RHE (for sequences see panel a) were used to drive *RFP* expression. The RFP signal is strongly reduced when expression is controlled by the mutated *EXPA7* promoter compared to WT. Red, RFP signal. **e,** Reduction of RFP signal intensity in *pEXPA7:RFP*/*pRD26:RD26-GFP7*/*rd26-1* seedlings compared to *pEXPA7:RFP*/*rd26-1* seedlings. **f,** Weak RFP signal in the root epidermis of *pEXPA7:RFP*/*pRD26:RD26-GFP7*/*rd26-1* seedlings. For e and f, six-day-old agar-grown seedlings were incubated in liquid ½ MS medium containing 15 µM ABA for 12 h. Cyan: RD26-GFP; magenta: RFP.

The promoter of the root hair-specific *EXPANSINA7* (*EXPA7*) gene harbours an RHE −132 bp upstream of the translation start codon (**Supplementary Fig. 6**). Considering that RHE might be an RD26 BS, we performed a yeast one-hybrid assay utilising a 500-bp upstream region of the *EXPA7* gene that harbours RHE. RD26 activated the wild-type *EXPA7* promoter in yeast, indicating physical interaction between RD26 and RHE; this activation of the *EXPA7* promoter was lost when nucleotides shared by RHE and the RD26 BS were mutated (**Fig. 5b**). Next, we performed an electrophoretic mobility shift assay (EMSA) employing a 40-bp *EXPA7* promoter fragment encompassing RHE. RD26 interacted with the wild-type promoter fragment, while this interaction was lost when core RHE nucleotides were mutated (**Fig. 5c**). Employing *pRD26:RD26-GFP*/*rd26-1* plants (28) we next tested binding of RD26 to the *EXPA7* promoter *in planta* by chromatin immunoprecipitation – quantitative polymerase chain reaction (ChIP-qPCR). Expression of *RD26* was induced by treating plants with ABA. The *EXPA7* promoter fragment encompassing the RD26 BS was significantly enriched in the precipitated chromatin revealing it as direct transcriptional target of RD26 *in planta* (**Fig. 5d**).

To corroborate the regulatory interaction between RD26 and the *EXPA7* promoter and its RHE, we employed the same wild-type and mutated versions of the *EXPA7* promoter to control expression of the *RED FLUORESCENCE PROTEIN* (*RFP*) reporter in transgenic Arabidopsis plants. As expected, we observed a strong RFP signal in root hair files of Col-0 plants transformed with the wild-type *EXPA7* promoter driving *RFP* expression (**Fig. 5e; Supplementary Video 2**); signal intensity dropped strongly when *RFP* expression was controlled by the mutated *EXPA7* promoter lacking RHE (**Fig. 5e; Supplementary Video 2**). We also expressed *RFP* driven by the wild-type *EXPA7* promoter in *rd26-1* and *pRD26:RD26-GFP*/*rd26-1* genetic backgrounds. Plants were treated with ABA to induce *RD26-GFP*, leading to a reduction of RFP signal in *pRD26:RD26-GFP*/*rd26-*1 compared to *RFP* expressed from the *EXPA7* promoter in *rd26-1* (**Fig. 5f**). As shown in **Figure 5f** and **Supplementary Video 3**, the RFP signal in the root epidermis strongly declined in the presence of RD26-GFP expression (i.e., high GFP signal). This result clearly demonstrates that RD26 represses the expression of *EXPA7*.

### RD26 directly suppresses *RSL4*, a key regulator of root hair development

To investigate how RD26 might control root hair development, we reanalysed our previously reported whole-seedling microarray data obtained from *RD26-IOE* seedlings induced with EST (28) for root hair-related genes affected by RD26. As we show below, RD26 lowers the expression of multiple root hair morphogenesis genes, as well as *RSL4*, a key positive transcriptional regulator of root hair development (9) (**Supplementary Fig. 7**). We confirmed downregulation of *RSL4* after EST treatment by qRT-PCR in independent biological *RD26-IOE* samples (**Fig. 6a**). Next, we crossed *RD26Ox* plants with plants expressing an RSL4-GFP fusion from the *RSL4* promoter (*pRSL4:RSL4-GFP*). In the absence of *RD26* overexpression, RSL4-GFP gave a strong GFP signal in nuclei of root hair cells (**Fig. 6b**) (9); this signal was strongly reduced in offspring of the cross, supporting the model that RD26 suppresses *RSL4* expression (**Fig. 6b; Supplementary Video 4**).

**Figure 6.**
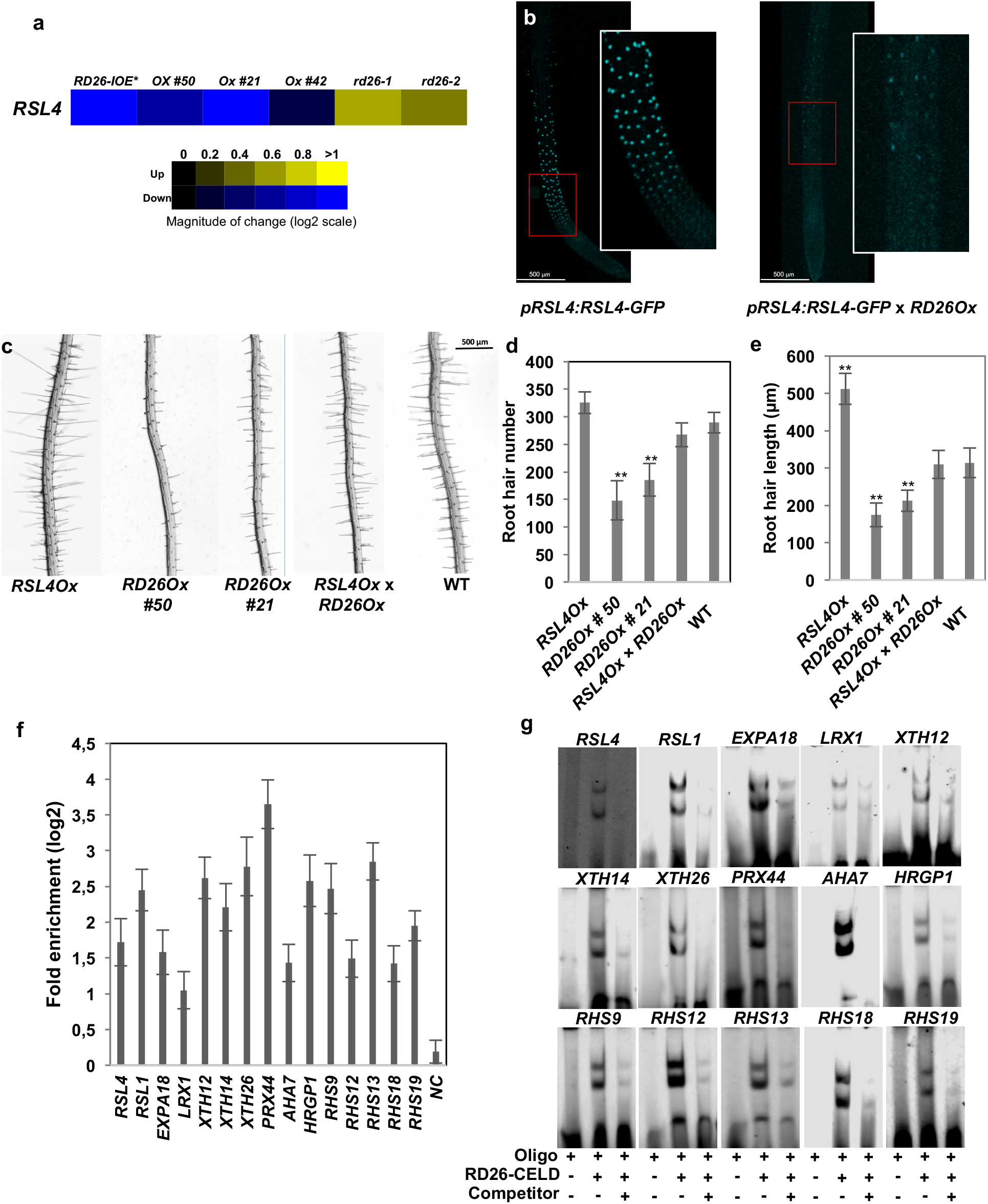
RD26 directly regulates genes important for root hair growth, including *RSL4*. **a,** Expression of *RSL4* is reduced upon *RD26* overexpression. Relative transcript abundance of *RSL4* (qRT-PCR). Column 1: expression ratios of genes in 2-week-old *RD26-IOE* seedlings after 5 h EST (10 µM) treatment compared to mock-treated plants. Columns 2 – 6: expression ratios of genes in 2-week-old seedlings of the indicated lines compared to WT. Data are the average of two biological replicates; each one includes three technical replicates. The heat map indicates expression ratios (log2): yellow, increased expression (between 0 and + 1 compared to control); blue, reduced expression (between 0 and - 1 compared to control). **b,** Fluorescence image of RSL4-GFP in root hair cell files of *pRSL4:RSL4-GFP* seedlings and the marker-resistant F2 progenies of the *pRSL4:RSL4-GFP x RD26Ox* cross. Red-boxed areas are enlarged on the right. **c,** The marker-resistant F2 progenies of the *RD26Ox x pRSL4:RSL4-GFP* cross exhibit a wild-type root hair phenotype. **d,** Root hair number. **e,** Length of root hairs in the first 10 mm of the root tip. Data are the means of 20 seedlings of two biological growth experiments ± SD. Asterisks indicate statistically significant differences compared with WT (ANOVA, ** *P* < 0.01). **f,** ChIP-qPCR. *RD26:RD26-GFP*/*rd26-1* and *rd26-1* control seedlings grown on ½ MS agar plates were transferred to ½ MS liquid culture and treated with 15 µM ABA, 24 h. Enrichment of promoter regions was quantified using qPCR. As a negative control (*NC*), qPCR was performed on a promoter (*At2g22180*) lacking RD26 binding sites. Means ± SD (two biological replicates, three technical replicates each). **g,** EMSA. RD26-CELD protein binds specifically to RD26 binding sites of promoters of target genes. Binding reactions were performed using ∼40-bp fragments of the promoters containing the RD26 binding site. RD26-CELD binds to labelled double-stranded oligonucleotides (middle lanes), while binding is not detected in the absence of RD26-CELD (left lanes) or when non-labelled competitor is added (100-fold molar excess; right lanes).

As RD26 downregulates *RSL4* expression leading to inhibition of root hair growth, we hypothesized that constitutive overexpression of *RSL4* in the *RD26Ox* background would reverse the negative effect on hair growth exerted by RD26. To test this, we crossed *RSL4* overexpressor (*RSL4Ox*) with *RD26Ox* plants. Root hair number and length increased by 13% and 63%, respectively, in *RSL4Ox* seedlings compared to WT, as expected (9). In contrast, root hair number and length were about 40% lower in *RD26Ox* seedlings than WT (**Fig. 6c-e**). The inhibitory effect of RD26 was completely lost in *RSL4Ox* × *RD26Ox* seedlings, indicating that *RSL4* expressed from the constitutive promoter (CaMV 35S) escapes the control by RD26.

An RD26 BS is present in the promoter of *RSL4* (1,632 bp upstream the translational start codon; **Supplementary Fig. 6**). We tested binding of RD26 to the *RSL4* promoter *in planta* by ChIP-qPCR using *pRD26:RD26-GFP*/*rd26-1* (28) (**Fig. 6f**). Expression of *RD26* was induced by treating plants with ABA. The *RLS4* promoter fragment encompassing the RD26 BS was significantly enriched in the precipitated chromatin revealing it as direct transcriptional target of RD26 *in planta* (**Fig. 6f**). EMSA experiments confirmed binding of RD26 to the respective promoter fragment of *RSL4* containing the RD26 BS (**Fig. 6g**).

Eighty-three genes are upregulated in *RSL4Ox* seedlings (9). Consistent with the model that RD26 is a negative regulator of *RSL4*, 37 of the 83 RSL4-induced genes are repressed in *RD26-IOE* and *RD26Ox* seedlings (**Fig. 7**). Moreover, 15 of the 18 root hair-specific genes induced by RSL4 are repressed by RD26.

**Figure 7.**
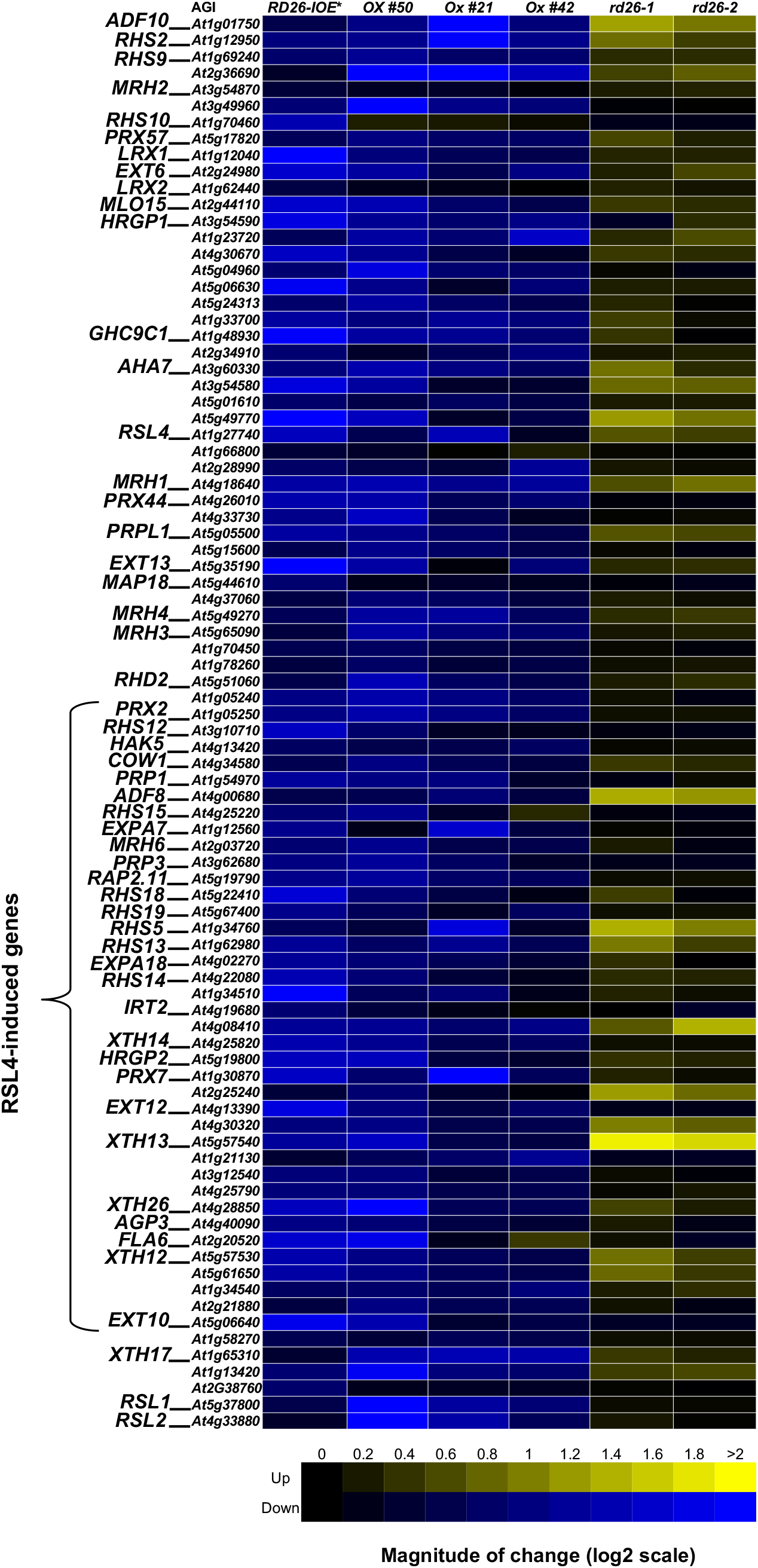
Root- and root hair-related genes of the RD26 regulon. Relative transcript abundance of the genes was determined by qRT-PCR. Along the x-axis, column 1 indicates the expression ratios of genes in two-week-old *RD26-IOE* seedlings after 5 h EST (10 µM) treatment compared to mock-treated plants. Columns 2 - 6 present the expression ratios of genes in two-week-old seedlings of the indicated lines compared to WT. Data are the average of two biological replicates; each one includes three technical replicates. The heat map indicates expression ratios (log2): yellow, increased expression (between 0 and + 2 compared to control); blue, reduced expression (between 0 and - 2 compared to control).

qRT-PCR analysis showed that expression of *RSL1* and *RSL2* was also significantly reduced in *RD26-IOE* seedlings after 5 h EST treatment compared to mock treatment, and the three *RD26ox* lines compared to WT (**Fig. 7**). The *RSL1* promoter harbours an RD26 BS (**Supplementary Fig. 6**). ChIP-qPCR and EMSA revealed binding of RD26 to the *RSL1* promoter, indicating it as a direct target of RD26 (**Fig. 6f, g**). However, we did not observe an effect of RD26 on the expression of *RHD6* and *RSL3*, or on the expression of the core regulatory TFs which are responsible for fate determination of hair and non-hair cells (**Supplementary Fig. 8**).

### RD26 suppresses root hair morphogenesis genes

Our analysis of the expression of root- and root hair-related genes in the whole-seedling microarray data obtained from *RD26-IOE* lines (28) identified 49 genes, including *RSL4*, with an at least 2-fold reduced expression 5 h after EST induction, compared to mock-treated samples (**Supplementary Fig. 7a** and **Supplementary Table 1**); qRT-PCR analysis confirmed reduced expression of all 49 genes in two new biological replications (**Fig. 7, Supplementary Table 2**). According to Genevestigator, most of the 49 genes are predominantly expressed in roots, and particularly in the root elongation zone and root hair cells (**Supplementary Fig. 9**), and 23 of these genes are amongst the 50 genes most up-regulated in root hair cells compared to non-root hair cells (**Supplementary Table 1**) (33). Of note, many of the 49 genes repressed by RD26 have previously been reported to play a role in root hair growth (see references in **Supplementary Table 1**).

We also analysed the microarray data for down-regulated root hair-related transcripts that might not have passed the 2-fold downregulation threshold, which resulted in 41 additional genes (**Supplementary Fig. 7b**, **Supplementary Table 1**); all genes were independently confirmed by qRT-PCR to be repressed by RD26 in *RD26-IOE* seedlings after 5 h EST treatment (**Fig. 7, Supplementary Table 2**). Furthermore, expression of the 86 (45 + 41) genes in two-week-old seedlings of three *RD26ox* and both *rd26* mutants revealed that 80 genes were repressed in *RD26ox* lines and 19 genes were induced by at least 1.5-fold in *rd26* mutants (**Fig. 7, Supplementary Table 2**). Won *et al*. (5) identified 31 root hair cell-specific genes, all of which possess an RHE in their promoters and are repressed in root hair-defective mutants; 21 of the 31 genes are repressed by RD26 (**Supplementary Table 1**).

### Single-cell RNA-seq confirms negative correlation between *RD26* and target genes

Single-cell RNA sequencing (scRNA-seq) is a recently established technology that provides transcriptome information of individual cells whereby patterns of gene expression are discovered by gene cluster analysis (34). Recently, scRNA-seq was employed to determine gene expression profiles in Arabidopsis roots (35, 36). We interrogated expression of *RD26* and its 86 repressed genes in the eight root-epidermis-related clusters reported by Ryu *et al*. (2019) (36). In accordance with our data, expression of *RD26* was considerably higher in N cells than H cells, while most of its target genes were higher expressed in H cells than N cells (**Supplementary Fig. 10a**). A correlation analysis over all eight root epidermis-related clusters showed negative correlations between the expression of *RD26* and its targets, while targets which are mainly involved in root hair outgrowth have positive correlations with each other (**Supplementary Fig. 10b; Supplementary Table 3**).

### RD26 represses multiple root hair morphogenesis genes by direct promoter binding

We found that in addition to *EXPA7*, *RSL1* and *RSL4*, putative RD26 BSs are present in the promoters of multiple other root hair-related genes repressed by RD26 (**Supplementary Fig. 6**). ChIP-qPCR demonstrated binding of RD26 *in planta* to 13 of them after ABA treatment, revealing them as direct targets (**Fig. 6f**); EMSA confirmed binding of RD26 to all 13 genes *in vitro* (**Fig. 6g**). Besides *EXPA7*, also *EXPA18* is specifically expressed in root hair cells; RD26 downregulates *EXPA18* (**Fig. 7**) by binding to its promoter as revealed by ChIP-qPCR and EMSA (**Fig. 6f, g**).

The formation of root hairs requires the action of diverse cell wall-modifying proteins including extensins (EXTs), arabinogalactan-proteins (AGPs), and proline-rich proteins (PRPs) (collectively called hydroxyproline-rich glycoproteins, HRGPs), as well as expansins (EXPAs) (37, 38). We found that RD26 represses various *HRGPs* and *EXPAs*, most of which play a role in root hair cell wall specification and expansion (**Supplementary Table 1**), including *LEUCINE-RICH REPEAT1* (*LRX1*; **Fig. 8**) (39, 40). The *LRX1* promoter harbours an RD26 BS (**Supplementary Fig. 6**), and ChIP-qPCR and EMSA confirmed RD26 as a direct upstream regulator (**Fig. 6f, g**). Similarly, RD26 represses *HRGP1* (*EXT2*) by binding to its promoter (**Fig. 6f, g; Fig. 7**). Several other *EXT* genes (*LRX2, EXT6*, *EXT10*, *EXT12*, *EXT13, At1g23720*, *At3g54580*, and *At4g08410*) are also repressed by RD26 (**Fig. 7; Supplementary Table 1**).

**Figure 8.**
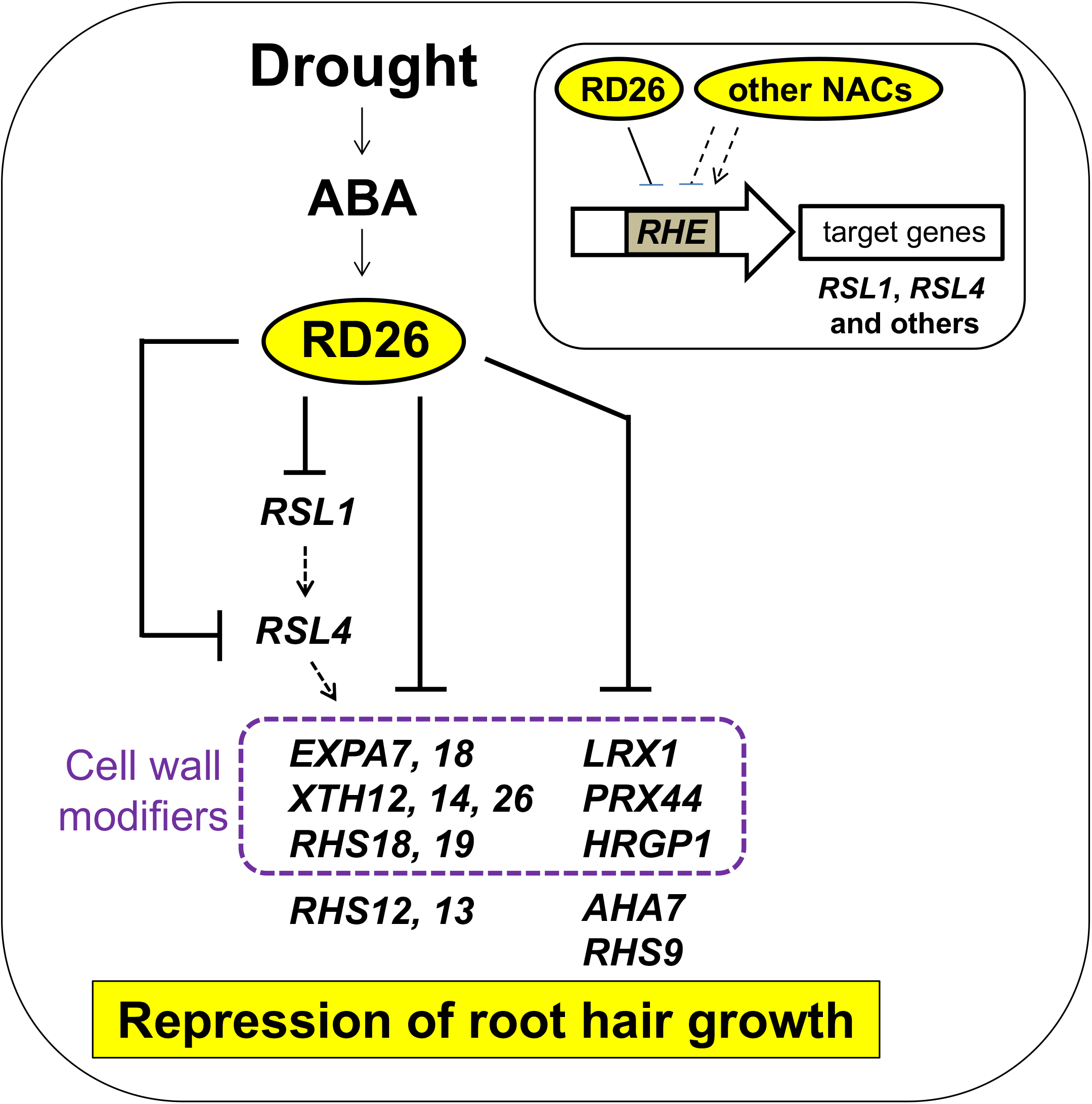
Model of the role of RD26 in suppressing root hair growth. Expression of NAC transcription factor gene *RD26* is induced by drought and the stress-triggered phytohormone abscisic acid (ABA). RD26 binds to the root hair-specific *cis*-element (RHE; inset) and directly suppresses expression of *RSL1* and *RSL4*, two bHLH transcription factors positively regulating root hair growth in Arabidopsis trichoblasts. In addition, RD26 directly suppresses the expression of several other root hair-specific genes mainly involved in cell wall modification by binding to the RHE *cis*-regulatory element present in their promoters, thereby contributing to the forward loop that limits root hair growth. This mode of action constitutes a robust gene regulatory network to repress root hair elongation under drought stress conditions. Besides RD26, additional but currently unknown NAC transcription factors (positively or negatively acting) might bind to RHE to control the expression of target genes (inset). Solid lines indicate direct interaction, while dashed lines indicate interaction of an unknown mode of action. Lines with an arrow head indicate positive regulation, while T-ending lines indicate negative regulation.

The *PRP* gene family in Arabidopsis encompasses three canonical members with *PRP1* and *PRP3* being expressed in a root hair-specific manner. Similarly, *PRPL1* is highly expressed in trichoblasts to trigger root hair growth (41). All three genes are transcriptionally repressed by RD26 (**Fig. 7**); in EMSA experiments, RD26 binds to BSs in the promoters of *PRP1* and *PRPL1* (**Supplementary Fig. 6** and **Supplementary Fig. 11**), suggesting they are RD26 targets. RD26 also represses the *AGP* genes *AGP3* and *FASCICLIN-LIKE ARABINOGALACTAN 6* (*FLA6*; **Fig. 7**), which are members of a gene regulatory network of root hair differentiation (10) and are induced by RSL4 (9).

Xyloglucan endotransglucosylases/hydrolases (XTHs) act on xyloglucans which are dominant cell wall hemicelluloses that interconnect cellulose microfibrils and thereby affect cell wall properties and cell expansion. XTHs have xyloglucan endohydrolase (XEH) or xyloglucan endotransglucosylase (XET) activity, or both, to cleave and reconnect xyloglucan crosslinks. The initiation of root hairs involves a boost of XET activity (15). Arabidopsis has 33 *XTH* genes, of which ten are specifically expressed in roots (15, 42, 43); RD26 represses six of them, *XTH12*, *13*, *14*, *17*, *19* and *26*, all of which encode enzymes with only XET activity (**Fig. 7; Supplementary Table 1**). Under non-stress conditions, these genes are specifically expressed in root hair cells (*XTH12*, *13*, *14* and *26*), or more generally in root cells (*XTH17* and *19*) (44, 45). RD26 BS are present in the *XTH12*, *13*, *14* and *26* promoters (**Supplementary Fig. 6**), and RD26 interacts with them *in planta* and *in vitro* (**Fig. 6f, g; Supplementary Fig. 11**).

Class III peroxidases are involved in the cleavage and crosslinking of cell wall polysaccharides during wall remodelling (37, 46–48). Nine class III *PEROXIDASES* expressed in root hair cells (*PRX7*, *PRX44*, *PRX57*, *PRX2*, *RHS18*, *RHS19*, *At1g34510*, *At3g49960* and *At1g05240*; (5, 10, 13, 26, 40, 49), are repressed by RD26 (**Fig. 7; Supplementary Table 1**), and root hair growth is impaired in *prx7*, *prx44* and *prx57* mutants (13, 49). ChIP-qPCR and EMSA revealed direct binding of RD26 to the *PRX44*, *RHS18* and *RHS19* promoters (**Fig. 6f, g; Supplementary Fig. 6**).

### Additional root hair-associated genes controlled by RD26

Several other genes involved in root formation are directly repressed by RD26, including *SEED AND ROOT HAIR PROTECTIVE PROTEIN* (*SRPP*)/*RHS13* (50) and the root hair-specific genes *RHS9*, *RHS12* and *RHS14* (5, 10) with currently unknown functions (**Fig. 6f, g; Fig. 7; Supplementary Fig. 11**). Additionally, *MORPHOGENESIS OF ROOT HAIR 1* (*MRH1*), a member of the leucine-rich-repeat (LRR) class of receptor-like kinases (10, 51), as well as *MRH2*, *MRH3*, *MRH4* and *MRH6* which all contribute to root hair morphogenesis (10, 51, 52) are repressed by *RD26* (**Fig. 7**).

Growth of root hairs requires cell wall acidification mediated by a plasma membrane proton-ATPase (PM H^+^-ATPase). Of the 12 PM H^+^-ATPase genes in Arabidopsis, only *AHA7* is positively involved in root hair development (53, 54). *AHA7* was the only gene significantly repressed (∼2-fold) by RD26, while its expression was induced in *rd26* mutants (1.5-fold; **Fig. 7**). The *AHA7* promoter harbours a putative RD26 BS (**Supplementary Fig. 6**), and ChIP-qPCR and EMSA revealed direct binding of RD26 to the *AHA7* promoter (**Fig. 7g, f**). RD26 also represses *ROOT HAIR DEFECTIVE2* (*RHD2*, *RbohC*) which encodes a respiratory burst oxidase homolog protein involved in the localized accumulation of reactive oxygen species and a subsequent Ca^2+^ influx into the cytoplasm needed for root hair growth (55–57) (**Fig. 7**). Similarly, it represses *CAN OF WORMS1* (*COW1*) encoding a phosphatidylinositol transfer protein which acts downstream of RHD2 (58, 59). RD26 also represses transcription factor RELATED TO AP2 11 (RAP2.11), a positive regulator of root hair growth, and its direct target *HIGH AFFINITY K^+^ TRANSPORTER 5* (*HAK5*) (9, 60) (**Fig. 7; Supplementary Fig. 11**).

### ABA differentially regulates genes of the RD26 root regulon

Given the fact that overexpression of *RD26* leads to root and root hair phenotypes reminiscent of those induced by ABA treatment, we analysed the ABA-dependent expression of the 86 RD26 root regulon genes. We observed that 63 genes were repressed by both ABA treatment and overexpression of *RD26* (**Supplementary Fig. 10**). The remaining genes are not regulated by ABA or they are affected by RD26 in a different direction (**Supplementary Fig. 12**).

To test whether repression of the 63 genes by ABA is further enhanced by *RD26* overexpression, we next analyzed the effect of ABA in *RD26-IOE* seedlings. Simultaneous treatment with ABA and EST further downregulated most of the 56 genes compared to the single treatments (EST or ABA). The 63 repressed genes fall into two groups, those that required RD26 for repression by ABA (group A; 34 genes), and those that were repressed by ABA largely independent of RD26 (group B; 29 genes; **Supplementary Fig. 12**). The repressive effect of ABA on group A genes disappeared or even changed to induction in *rd26* knockout mutants, clearly demonstrating that ABA requires RD26 for their suppression, while group B genes retained their ABA responsiveness in *rd26* knockout mutants. Interestingly, *RSL4* and many of its targets belong to group A. We, therefore, conclude that the repressive effect of RD26 on *RSL4* and its targets is likely the only pathway by which ABA represses root hair development (**Supplementary Fig. 12**).

Although the ABA-dependent repression of group B genes was further enhanced by EST-mediated induction of *RD26* expression in *RD26-IOE* seedlings (like in group A), ABA was still able to exert its effect in *rd26* mutants demonstrating that ABA represses the transcription of these genes *via* both, RD26-dependent and -independent pathways.

### The function of RD26 is evolutionary conserved across plant families

Plants of the orders Brassicales and Solanales exhibit the same type of position-dependent root hair patterning (4). To evaluate the effect of RD26 on root hairs in a representative species of the Solanales, we generated transgenic tomato (*Solanum lycopersicum*) plants expressing tomato *RD26* (*SlRD26*) from the EST-inducible promoter and found that induction of *SlRD26* suppresses root hair growth in tomato, as in Arabidopsis (**Supplementary Fig. 13a-c**). Furthermore, we found that SlRD26 represses genes orthologous to the key Arabidopsis root hair genes (**Supplementary Fig. 13d**) indicating an evolutionary conserved role of RD26 for controlling root hair growth in diverse plant species.

## Discussion

Root hair development is a plastic process controlled by environmental cues. While nutrient deficiency generally triggers root hair growth, drought and salinity stress typically decrease it (6, 9, 18). We show here that NAC transcription factor RD26 plays an important role in controlling root hair growth. We demonstrate that RD26 mainly suppresses genes operating in root hair morphogenesis and outgrowth rather than hair cell fate determination, a process controlled by core developmental TFs whereby TTG1, GL3, EGL3, WER and MYB23 govern non-hair fate specification, while CPC, TRY and ETC1 control hair cell fate specification (5, 10, 61–66). None of the core cell fate-determinant TFs is significantly affected by RD26 (**Supplementary Fig. 8**). In trichoblasts, *RHD6* and *RSL1* are induced by the core TFs, and RHD6 (and less so RSL1) induces several genes involved in root hair morphogenesis. RSL2 and RSL4 act downstream of RHD6 and RSL1 (8–11). RD26 and ABA do not affect *RHD6* expression, while the majority of the genes downstream of RHD6 and having a role in root hair outgrowth are repressed by both. Here, we show that RD26 directly represses both, *RSL1* and *RSL4*. Thus, during drought stress the ABA-triggered induction of *RD26* represses root hair outgrowth genes downstream of RHD6.

During polar trichoblast outgrowth, cell walls are modified to enhance tensile strength; the structural changes are mainly supported by the induction of *HRGP*, *EXP*, *XTH*, and *PRX* genes. Of note, 17 *HRGP* genes are repressed by RD26; a role in root hair cell wall specification and expansion has been reported for most of them (**Supplementary Table 1**). As to *EXP* genes, *EXPA7* and *EXPA18*, which both have a specific role in root hair cells, are directly repressed by RD26. Furthermore, of the ten *XTH* genes expressed in the Arabidopsis root (43), six are repressed by RD26 (*XTH12*, *13*, *14*, *17*, *19* and *26*). Notably, the initiation of root hair growth is accompanied by a boosted XET activity specifically at the position where a bulge forms for future root hair formation (15). Intriguingly, all six enzymes encoded by the RD26-repressed *XTH* genes only exhibit XET activity (44, 45); *XTH12*, *14* and 26 with a specific expression in trichoblasts/root hair cells are directly repressed by RD26. In addition, RD26 suppresses nine *PRX* genes typically expressed in root hair cells (5, 10, 13, 26, 40, 49).

An important further result of our study is that RD26 binds to the RHE *cis*-regulatory element required for the control of the transcription of root hair-expressed genes (4). It has previously been reported that RSL4 binds to RHE as a positive regulator of gene expression (67). In contrast, RD26 functions as a negative regulator of such genes during drought or osmotic stress, indicating that it competes for binding to RHE during drought stress thereby counteracting the root hair growth promoting function of RSH4. In addition, considering that RHE is bipartite in nature (4) and that it strongly resembles the binding site of NAC TFs (28, 68, 69) additional NAC factors might be involved in the control of the root hair morphogenic genes. Identifying them and testing their interplay with RD26 in regulating the expression of root hair-expressed genes will be an important endeavour in the future. Furthermore, transcription factor OBF BINDING PROTEIN4 (OBP4) suppresses root hair growth and *RSL2* expression in Arabidopsis (70), suggesting the involvement of multiple transcriptional regulators in controlling stress-dependent growth of root hairs.

An important further finding of our study is that the control of stress-dependent root hair development is conserved in other plant species, as we demonstrate here for the solanaceous crop tomato (**Supplementary Fig. 13**). This observation may be exploited in the future for fine-tuning root hair growth under abiotic stress conditions in other crop species as well.

To summarize, NAC transcription factor RD26 controls adaptive root hair development in response to drought and osmotic stress by controlling the expression of multiple key genes crucial for this process (**Fig. 8**). The action of RD26 maybe counteracted, or affected, by other NAC transcription factors not yet identified. RD26 is evolutionary conserved in monocot and dicot angiosperms, as well as in *Amborella trichopoda* (the most basal lineage of angiosperms), lycophytes (*Selaginella moellendorffii*), and liverworts (*Marchantia polymorpha*; **Supplementary Fig. 14**), suggesting that it may have played an important role in early land plant evolution. Studying the precise role of *RD26* orthologs in these species and their downstream gene regulatory networks will provide interesting insights into this evolutionary process which was critically important for the population of our planet.

## Materials and Methods

Experimental details including citations are provided in **SI Appendix.**

### Plant growth

Seedlings were grown in a climate chamber with 16 h day light provided by fluorescent light (∼100 µmol m^-2^ sec^-1^) and a day/night temperature of 20/16°C and a relative humidity (37) of 65%.

### cDNA synthesis and gene expression analysis

Transcriptome studies using Affymetrix ATH1 microarrays were reported in Kamranfar *et al.* (2018) (28). Expression data are available from the NCBI Gene Expression Omnibus (GEO) repository (www.ncbi.nlm.nih.gov/geo/) under accession number GSE100654.

### DNA-RD26 interaction studies

Recombinant RD26-CELD fusion protein was expressed in *Escherichia coli*, purified and used in electrophoretic mobility shift assays (EMSA) as described (28) using 5′-DY682-labeled oligonucleotides (**Supplementary Table 4**).

For chromatin immunoprecipitation – quantitative PCR (ChIP-qPCR), Arabidopsis *pRD26:RD26-GFP*/*rd26-1* and *rd26-1* (as control) seedlings were used. To induce *RD26* expression, eight-day-old seedlings grown on ½ MS agar medium were transferred to ½ MS liquid medium containing 15 µM ABA and harvested 24 h later. Anti-green fluorescent protein (anti-GFP) antibody was used in ChIP to immunoprecipitate protein–DNA complexes as previously described (71). To check binding of RD26 to the promoters of the target genes, qPCR was performed using primers amplifying the promoter regions in close vicinity of the RD26 BS (**Supplementary Table 4**). Primers annealing to promoter regions of a gene lacking an RD26 BS (*At2g22180*) were used as negative control. The ChIP experiment was run in two biological replications with three technical replications per assay.

### Root phenotyping

Seedlings kept vertically for 12 days were used for analysis. Images of the plates were captured using a HP flatbed scanner (HP C7716) and the length of PR and all visible LRs was quantified using ImageJ (https://imagej.nih.gov/ij/). Developmental stages of lateral root primordia were identified as described (72). A detailed description of the root hair phenotyping is provided in **SI Appendix, SI Materials and Methods.**

## Acknowledgements

We thank the University of Potsdam (UP) and the MPI of Molecular Plant Physiology (MPI-MP) for supporting our research. IK thanks the Iran Ministry of Science and Chamran University of Ahvaz/Iran for financial support. We thank Liam Dolan (University of Oxford) for providing Arabidopsis *RSL4* transgenic lines, Xinnian Dong (Duke University, USA) for providing the *triple nac* mutant seeds, and Malcolm Bennett (University of Nottingham, UK) for sharing experimental procedures on LR phenotyping. We thank Arun Sampathkumar and Mastoureh Sedaghatmehr (MPI-MP) for support in confocal microscopy and western blot analysis, respectively, Vikas Devkar (MPI-MP) for providing the *SlRD26* cDNA, Dagmar Kupper (UP) for generating the *SlRD26-IOE* construct, and Antje Schneider (UP) for tomato transformation.

## Funding

This research was supported through funding by the Deutsche Forschungsgemeinschaft (FOR 948; MU 1199/14–2, BA4769/1–2, and ERA-CAPS ‘AbioSe’, MU 1199/16-1).

### Author contributions

IK, SB and BM-R initiated the study; IK and BM-R designed the experiments which were performed by IK. IK and BM-R wrote the manuscript with contributions from SB. All authors agreed with the final manuscript and its submission for publication.

---

**Supplementary Figure 1.**
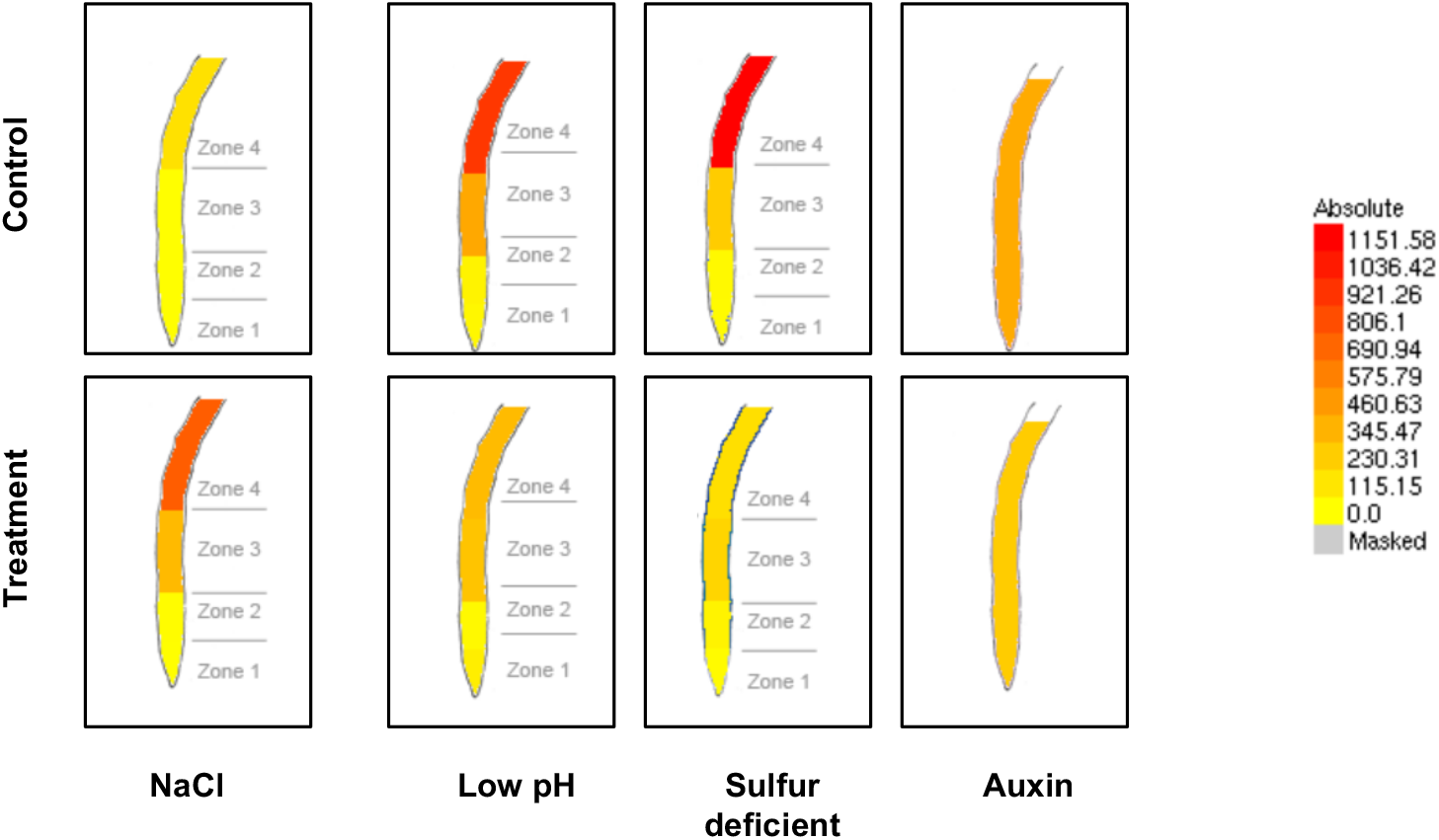
Expression of *RD26* in different segments of the Arabidopsis root. Induction of *RD26* by salt stress and repression by low pH, sulphur deficiency and auxin, mainly in the maturation zone of the root (‘Zone 4’) (Dinneny *et al*., 2008; Iyer-Pascuzzi *et al*., 2011; Bergmann *et al*., 2013). Data extracted from the eFP browser (https://bar.utoronto.ca/efp/cgi-bin/efpWeb.cgi).

**Supplementary Figure 2.**
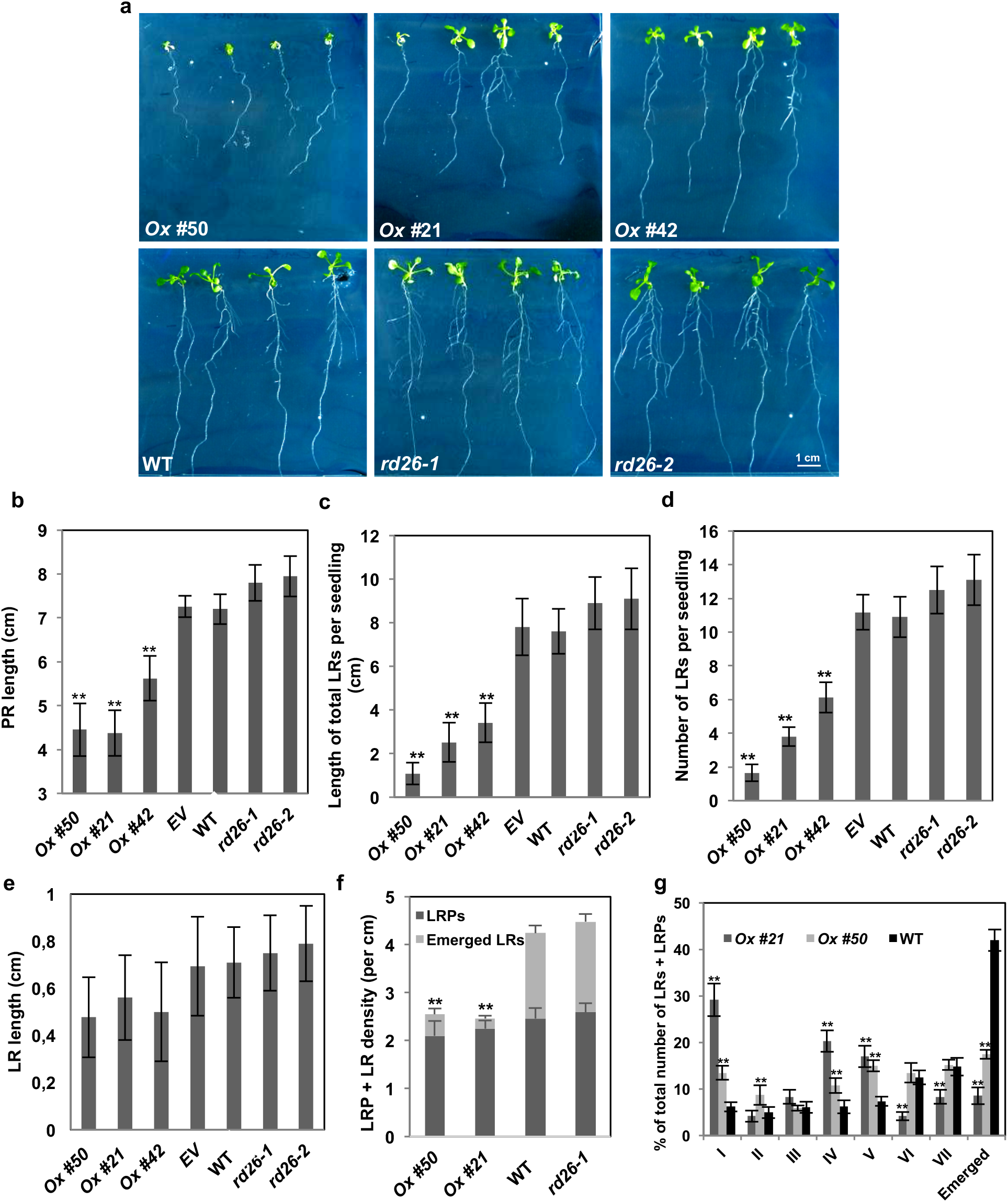
RD26 alters root-related growth parameters. **a,** Seedlings grown on **½** MS medium at 12 days after sowing. **b,** *RD26ox* lines have shorter primary roots (PR) and **c,** shorter total lateral roots (LRs) per seedling, which mainly resulted from reduced number of LRs and LRs + LR primordia (LRP) (see panels **d-f**; *n* = 20). **g,** Developmental stages of lateral root primordia in 12-day-old seedlings. Data are means obtained from three biological replicates ± SD. Asterisks indicate statistically significant differences form WT or *EV* (ANOVA, ** *P* < 0.01).

**Supplementary Figure 3.**
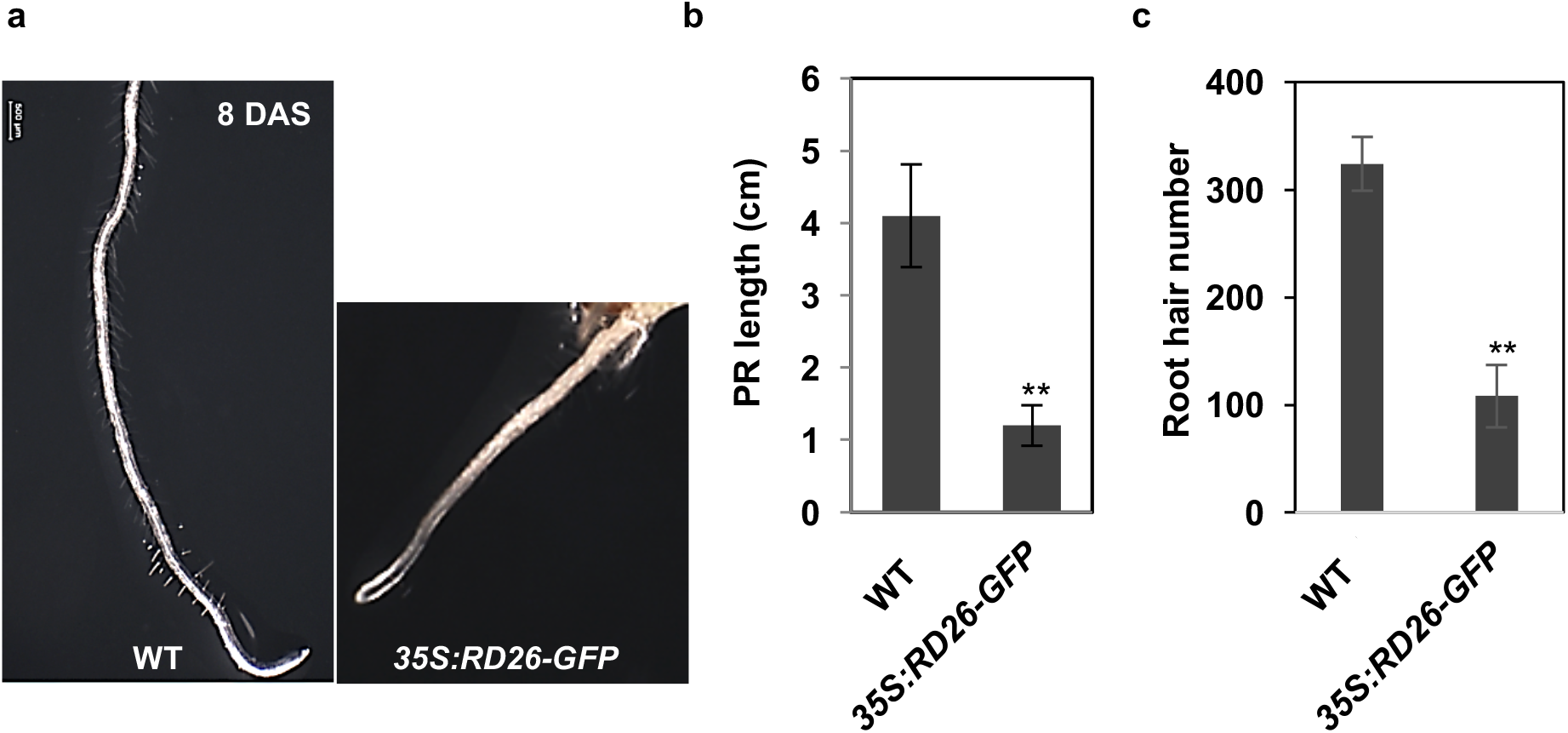
Constitutive overexpression of *RD26* represses primary root growth and root hair formation. **a,** MS-grown seedlings at 8 DAS. **b,** PR length and **c,** root hair number are reduced in *35S:RD26-GFP* seedlings compared to WT. The data shown are mean values obtained from 20 seedlings grown in three biological replicates ± SD. Asterisks indicate statistically significant differences from WT (Student’s *t*-test, ** *P* < 0.01).

**Supplementary Figure 4.**
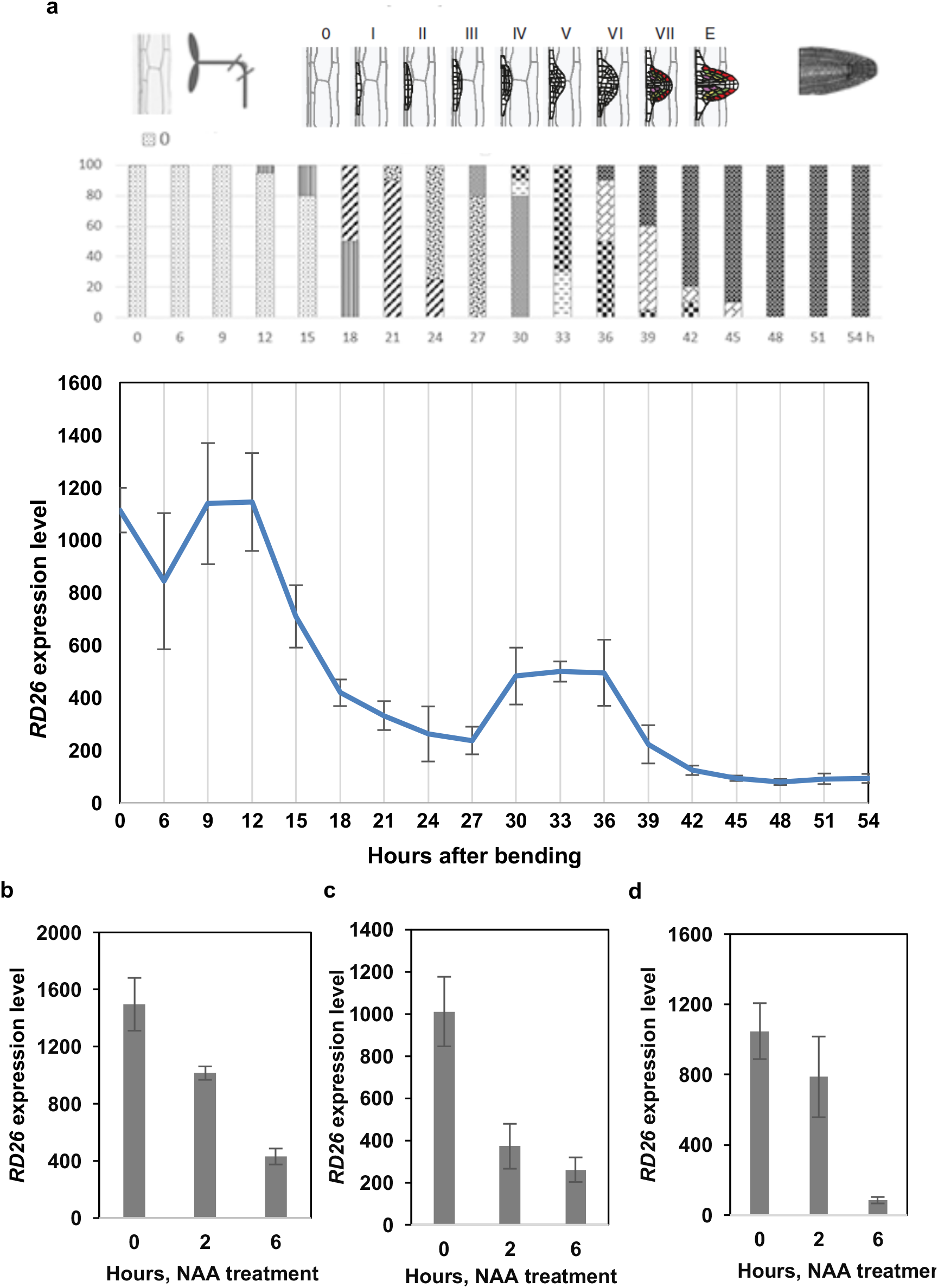
*RD26* is repressed during LR development. **a,** Repression of *RD26* expression during eight stages (I-E) of LR development. Data and the upper figure were obtained from the Arabidopsis eFP browser (http://bar.utoronto.ca/efp/cgi-bin/efpWeb.cgi). The original gene expression data were reported by Voß *et al*. (2015); LR induction was triggered by rotating vertically grown Col-0 seedlings by 90 degrees. Transcriptome profiles were analysed using Affymetrix ATH1 microarrays. **b, c,** *RD26* is repressed in roots of Arabidopsis Col-0 seedlings treated with 10 µM naphthaleneacetic acid (NAA) to synchrounsly induce LR initiation. Data in (b) and (c) were obtained from Vanneste *et al*. (2005) and De Rybel *et al*. (2012), respectively. **d,** *RD26* is repressed in xylem pole pericycle cells when LR initiation is induced by treatment with 10 µM NAA. Data were obtained from De Smet *et al*. (2008).

**Supplementary Figure 5.**
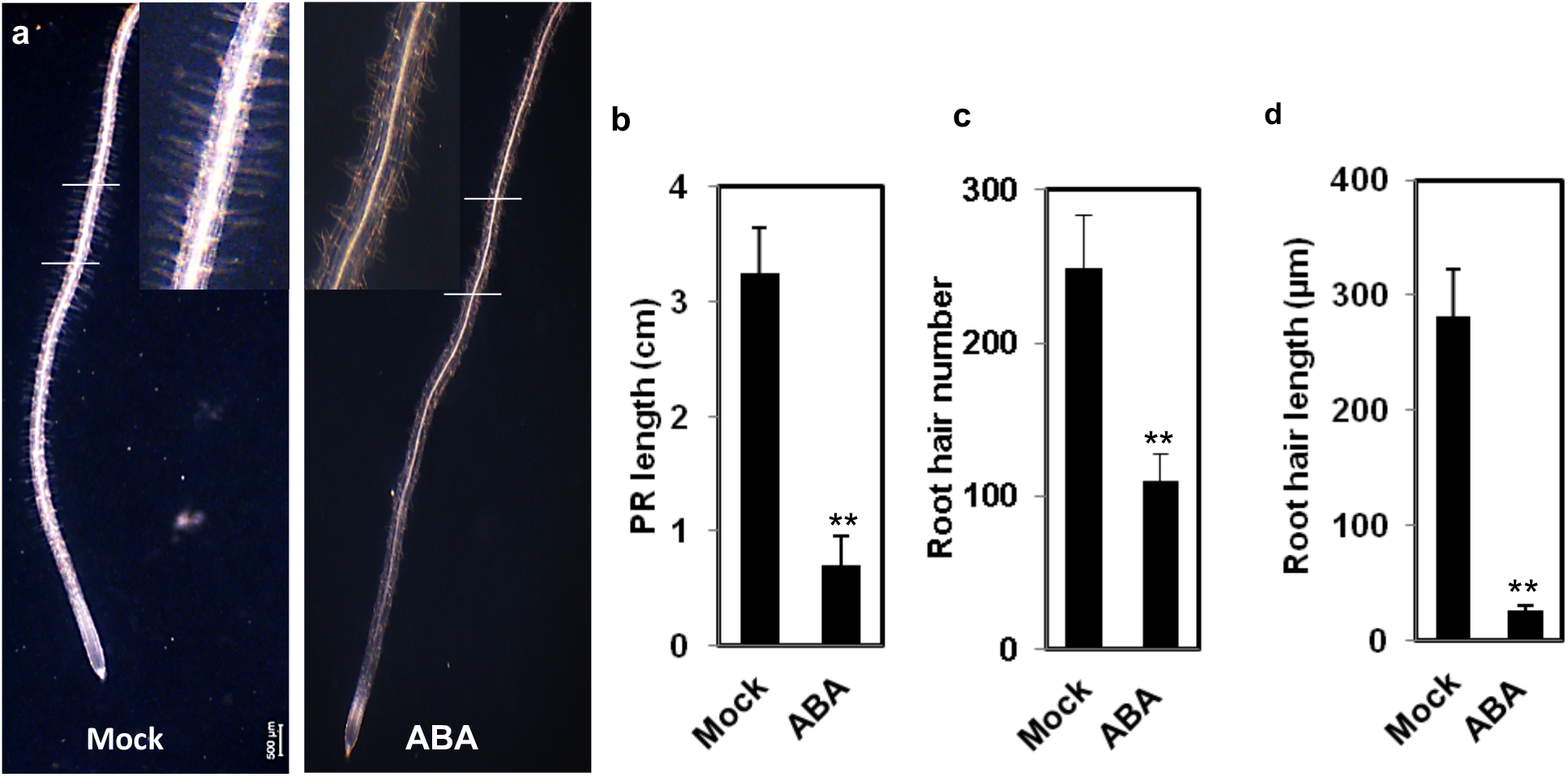
ABA represses root and root hair growth. **a,b,** ABA (15 µM) arrests primary root (PR) growth and **c,** reduces root hair number and **d,** length in Col-0 seedlings. Seedlings were transferred to ABA- or mock-containing plates at 5 DAS and roots were analyzed at 10 DAS (*n* = 20). Phenotyping of the root hairs was performed for the first 10 mm to the root tip. The data shown are mean values obtained from three biological replicates ± SD. Asterisks indicate statistically significant differences from mock-treated seedlings (Student’s *t*-test, ** *P* < 0.01).

**Supplementary Figure 6.**
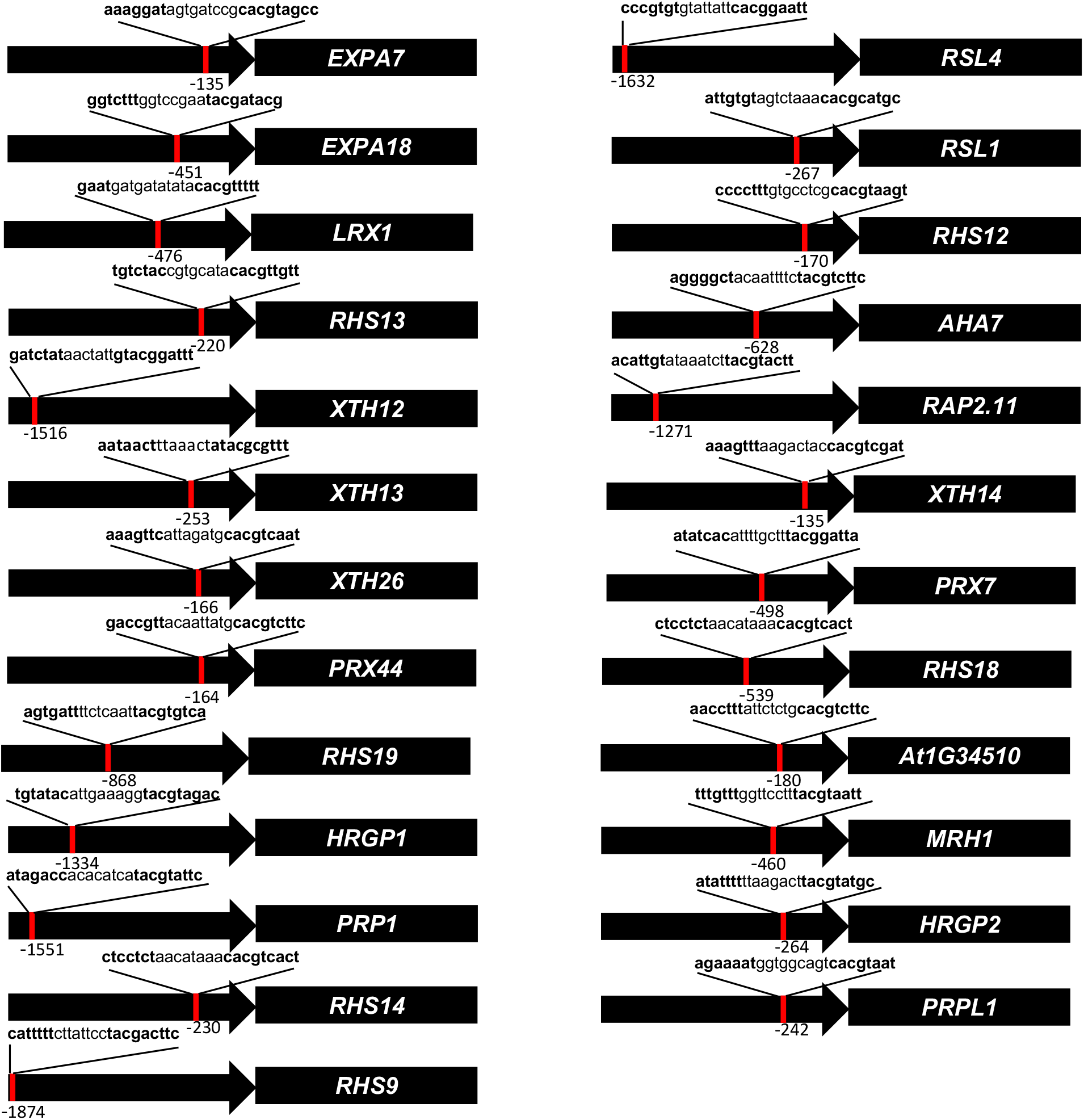
RD26 binding sites in the promoters of root hair-related target genes. Numbers indicate the 5’ position of the first nucleotide of the binding site upstream of the translation start codon (ATG). The RD26 binding site in the *EXPA7* promoter encompasses the RHE reported earlier (Kim *et al*., 2006).

**Supplementary Figure 7.**
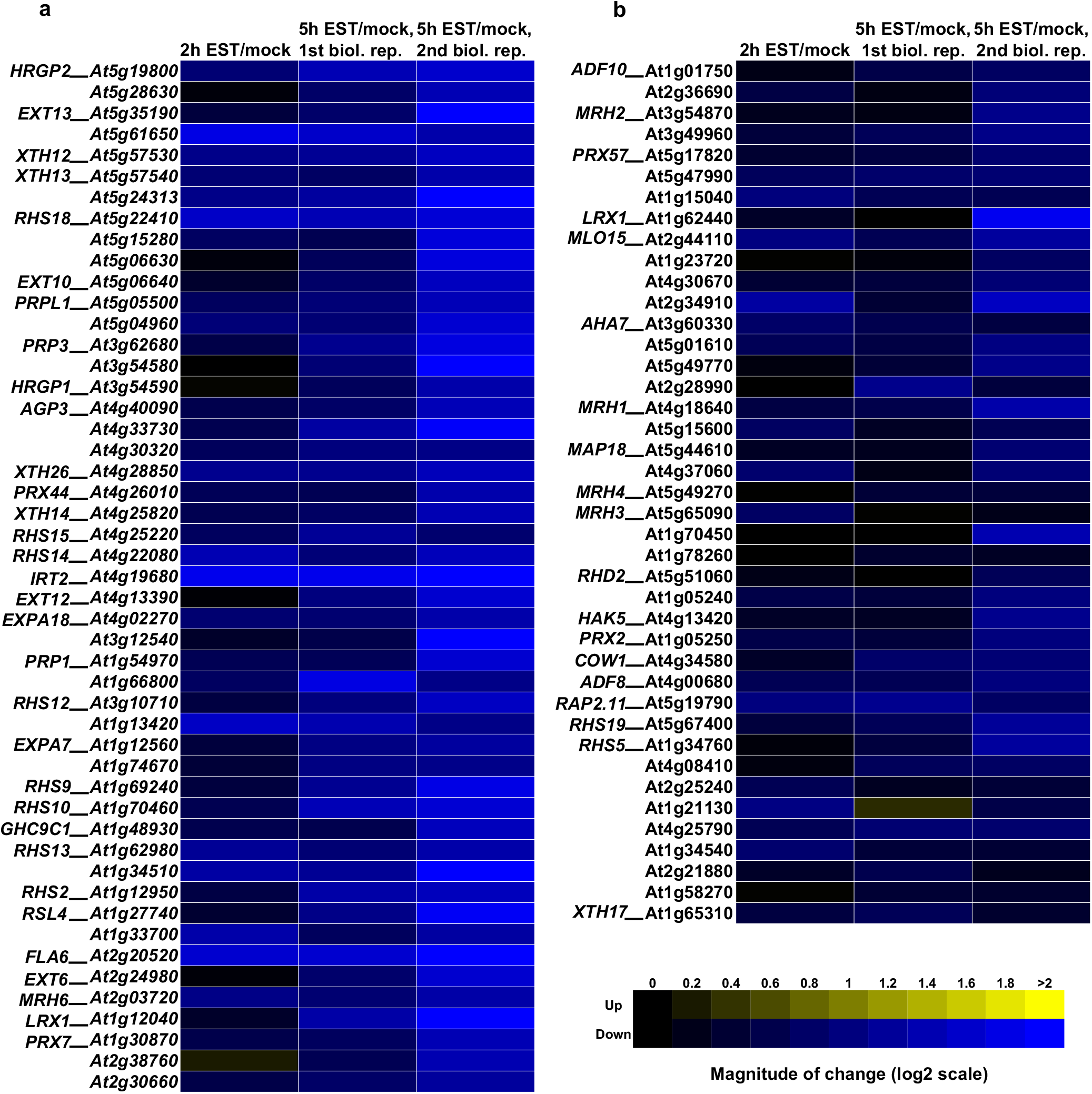
RD26 downregulated genes identified by Affymetrix ATH1 microarray analysis. Note, that the majority of the genes possess a function in root and root hair development. Relative transcript abundance of the genes was determined in EST-treated (10 µM EST) *RD26-IOE* plants compared to mock-treated controls (0.1% ethanol, v/v). Columns 2 and 3 represent data from two different biological replicates. The heat map indicates expression ratios (log2): yellow, increased expression (between 0 and + 2 compared to control); blue, reduced expression (between 0 and −2 compared to control).

**Supplementary Figure 8.**
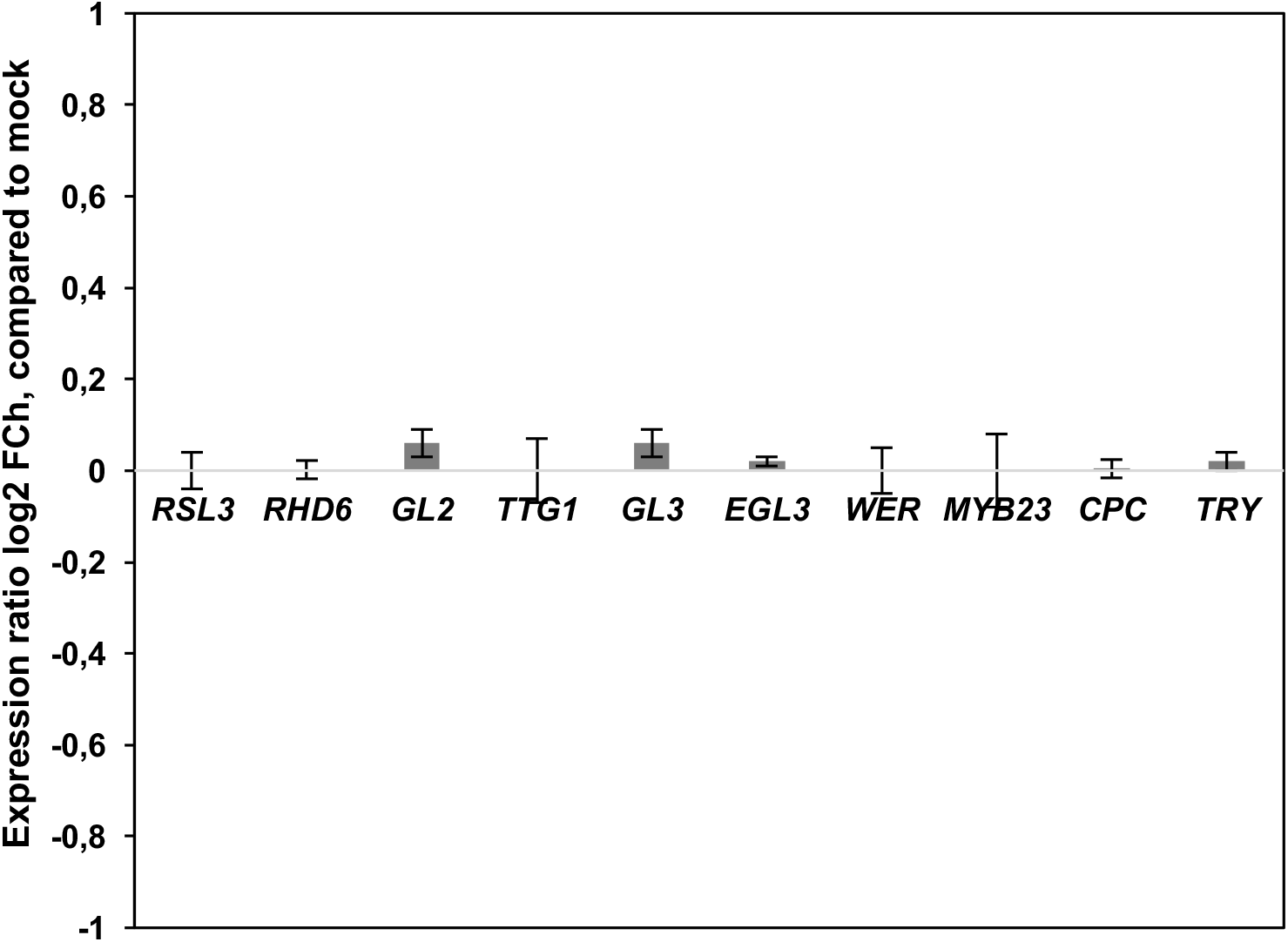
TFs involved in epidermis morphogenesis, but not regulated by RD26. Gene expression in 2-week-old, EST-treated *RD26-IOE* seedlings (compared to mock-treated seedlings, 5 h treatment) as determined by qRT-PCR. Numbers at the y-axis indicate log2 fold-change (FCh) expression ratio compared to the respective controls. Data are the means of three independent biological replicates ± SD.

**Supplementary Figure 9.**
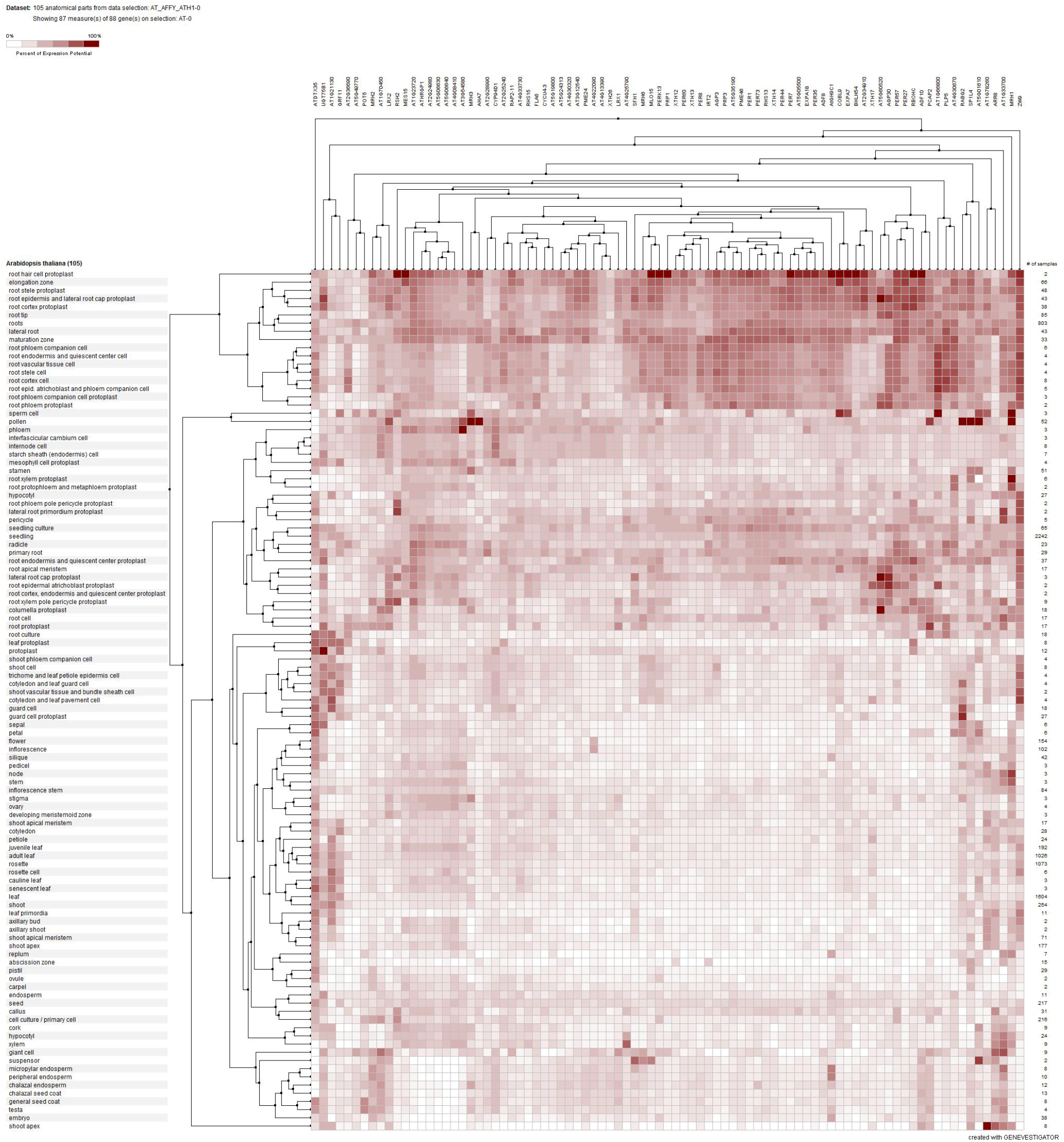
Tissue-specific expression profile of the 49 genes repressed by RD26. Hierarchical clustering was performed over genes and tissues using Euclidian distance method for available microarray data sets in Genevestigator tool (www.genevestigator.com). Darker color indicates higher gene expression.

**Supplementary Figure 10.**
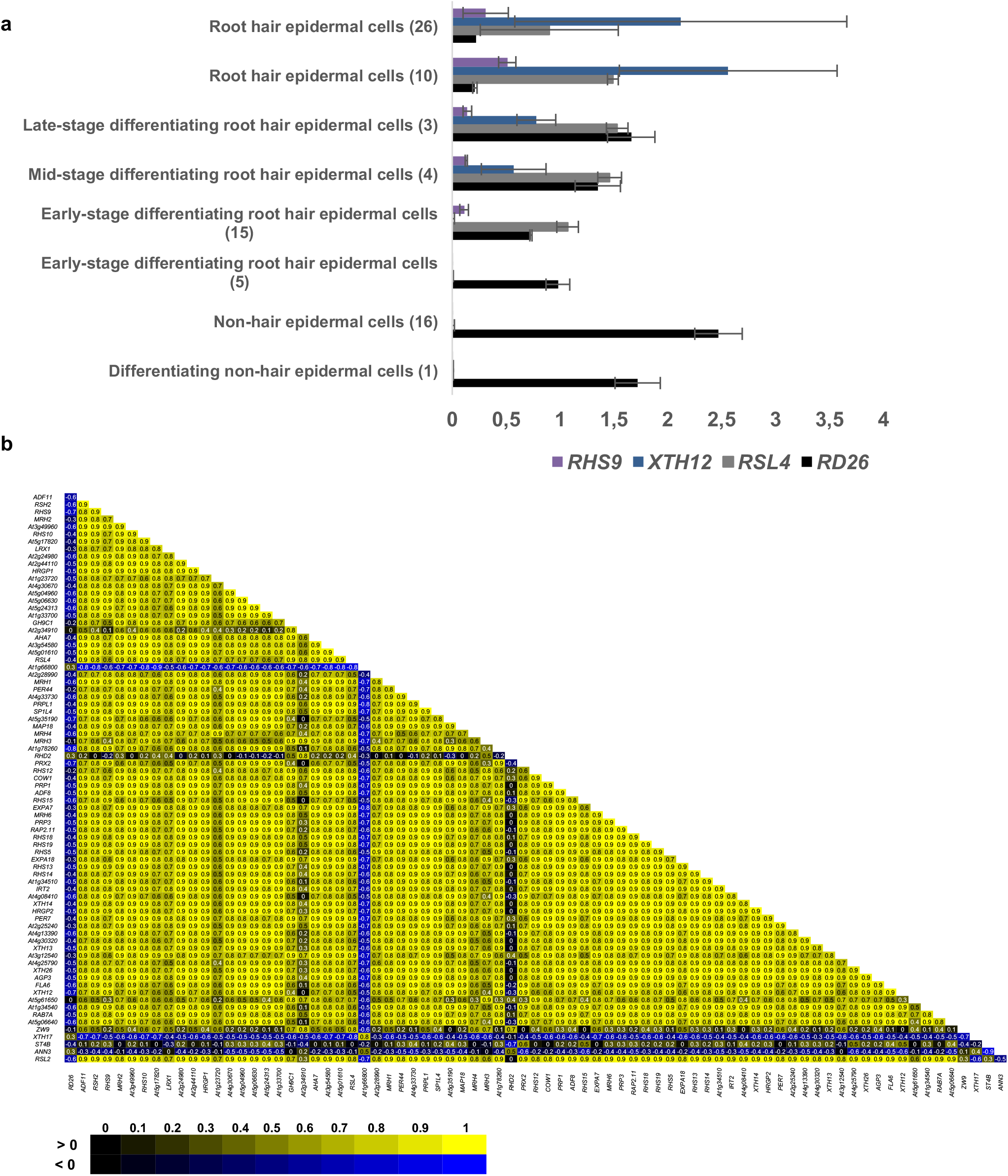
Single-cell RNA-seq supports a negative regulatory connection between *RD26* expression and expression of root hair morphogenesis genes. **a,** Expression of *RD26* and key root hair-related target genes in root epidermis clusters of single-cell RNA-seq data obtained from Ryu *et al*. (2019). Note, that *RD26* is higher expressed in N cells than H cells, in contrast to its repressed targets. **b,** Heatmap of Pearson correlation coefficients for all gene pairs (*RD26 vs*. its targets in the root hair regulon). While *RD26* expression is negatively correlated with the expression of its targets, the targets themselves mostly show a positive correlation with each other. The Pearson coefficient of correlation was calculated for the expression of each pair of genes over all eight epidermis cell clusters. Of the 86 genes of the RD26 root hair regulon, expression of 78 genes was detected and included in the analysis.

**Supplementary Figure 11.**
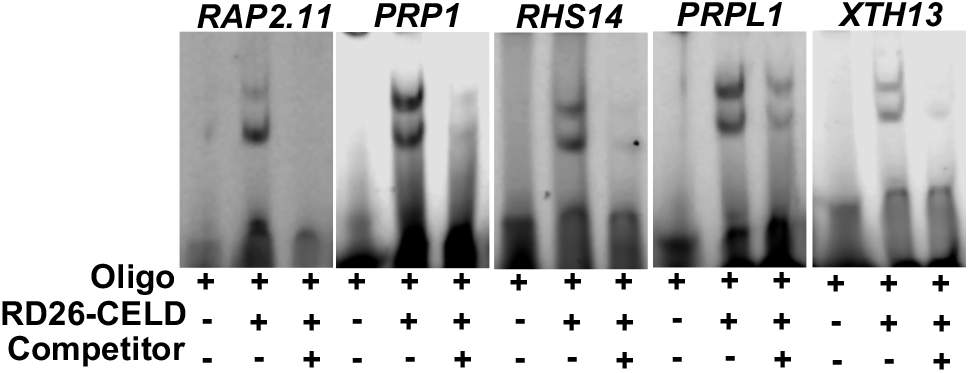
EMSA demonstrates binding of RD26 to promoters of root hair genes repressed by RD26. RD26-CELD protein binds specifically to RD26 binding sites of promoters of target genes. Binding reactions were performed using ∼40-bp fragments of the promoters containing the RD26 binding site. RD26-CELD binds to labelled double-stranded oligonucleotides (middle lanes), while binding is not detected in the absence of RD26-CELD (left lanes) or when non-labelled competitor is added (100-fold molar excess; right lanes).

**Supplementary Figure 12.**
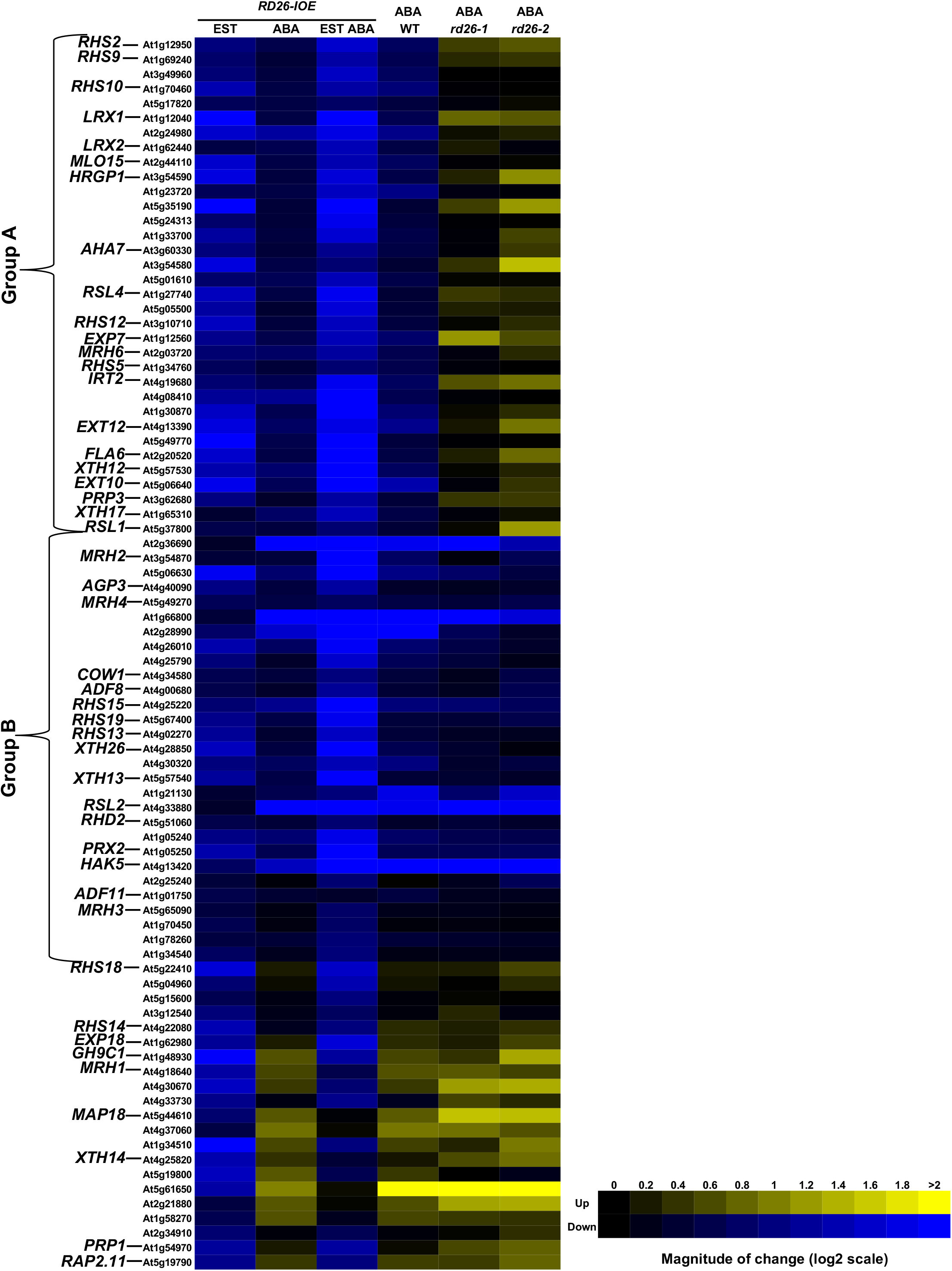
ABA differentially regulates genes of the RD26 root regulon. Root-related genes repressed by RD26 fall into groups based on the effect of ABA on their expression and the dependency on RD26. Genes in group A and B are down-regulated by both, RD26 and ABA. While in group A the ABA repression effect is dependent on RD26 (compare ABA effect on gene expression in *rd26* mutants and WT), ABA represses group B genes independently. Relative transcript abundance of the genes was determined using qRT-PCR. Along the x-axis columns 1 - 3 indicate the expression ratios of genes in two-week-old *RD26-IOE* seedlings after 5 h EST (10 µM), ABA (10 µM) or ABA + EST treatments compared to mock-treated plants. Columns 4 - 6 show the expression ratios of genes in two-week-old seedlings of indicated lines after 5 h ABA (10 µM) treatment compared to mock-treated plants. Data are the average of two biological replicates; each one includes three technical replicates. The heat map indicates expression ratios (log2): yellow, increased expression (between 0 and + 2 compared to control); blue, reduced expression (between 0 and −2 compared to control).

**Supplementary Figure 13.**
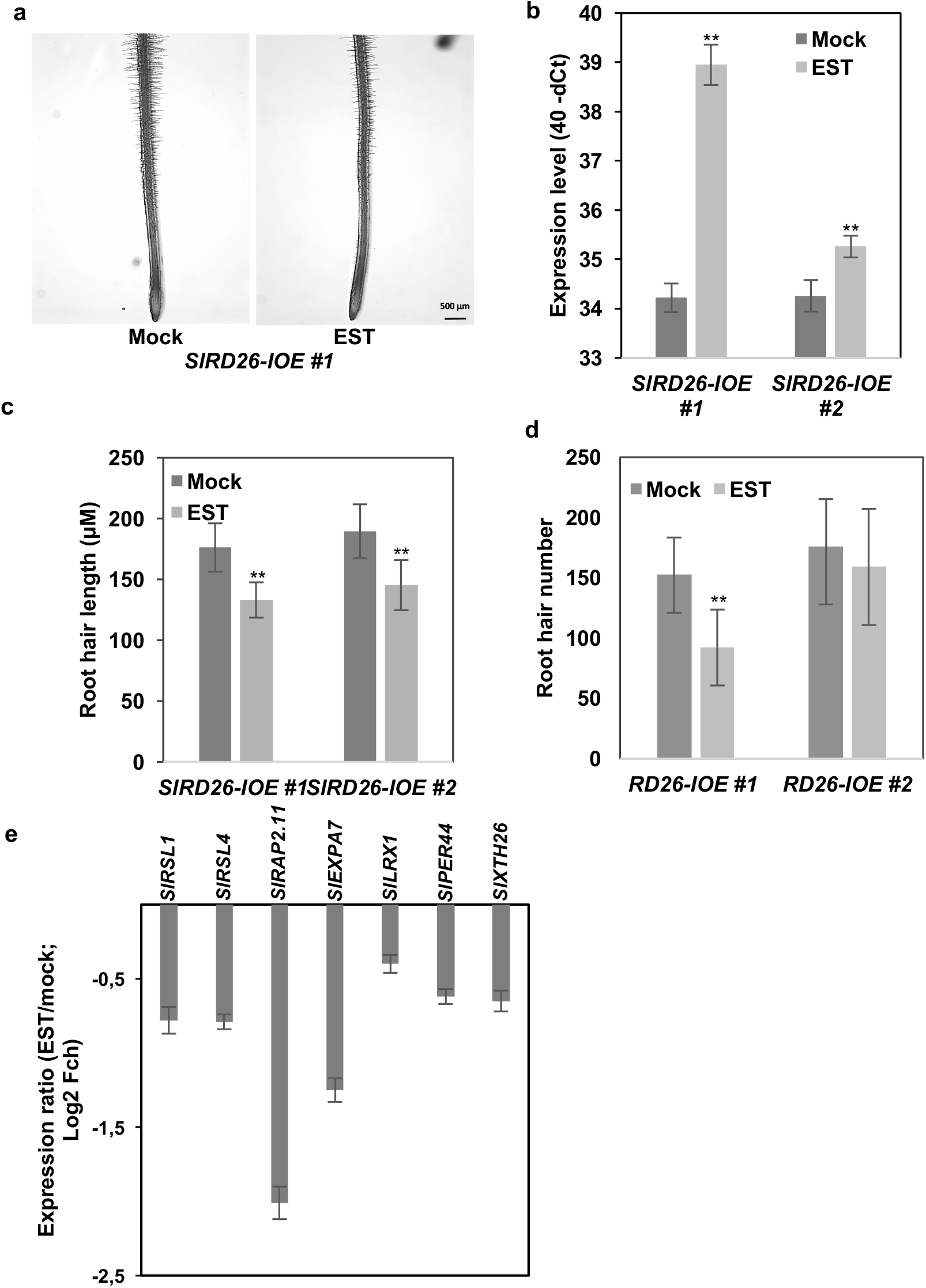
Induction of *SlRD26* represses root hair growth in tomato. **a,** Induction of *SlRD26* in *SlRD26* inducible overexpression plants (*SlRD26-IOE*) represses root hair growth. **b,** Root hair length. **c,** Root hair number. **d,** Induction of *SlRD26* represses orthologues of Arabidopsis root hair genes in *RD26-IOE* tomato plants. Seedlings were transferred to MS agar plates containing EST (15 µM) or ethanol (0.15% [v/v]; mock treatment). Phenotyping of root hairs was performed within the first 10 mm to the root tip at 4 DAT. Asterisks indicate statistically significant differences from mock-treated seedlings (ANOVA, ** *P* < 0.01). Tomato orthologues of the Arabidopsis genes have the following gene codes: *SlRD26*, *Solyc07g063410*; *SlRSL1*, *Solyc06g005000*; *SlRSL4*, *Solyc04g077960*; *SlRAP2.11*, *Solyc01g005630*; *SlEXPA7*, *Solyc08g080060*; *SlLRX1*, *Solyc11g005150*; *SlPRX44*, *Solyc10g084200*; *SlXTH26*, *Solyc12g007250*.

**Supplementary Figure 14.**
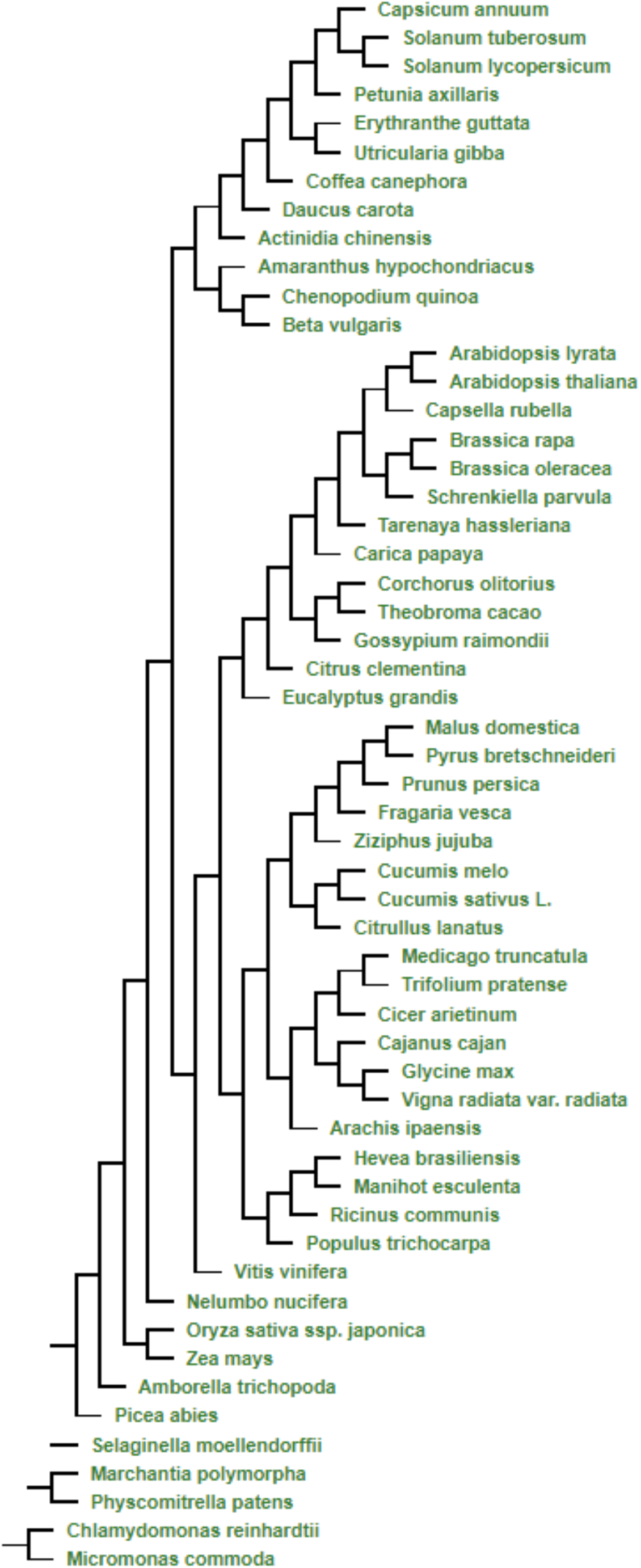
Phylogenetic tree of *RD26* orthologs in plants. Data were obtained from PLAZA 4.0 (https://bioinformatics.psb.ugent.be/plaza/; Van Bel *et al*., 2018).

## SI Materials and Methods

### General methods

Chemicals and kits were obtained from Sigma-Aldrich (Munich, Germany), Qiagen (Hilden, Germany), or Thermo Fisher Scientific (Berlin, Germany). Oligonucleotides were obtained from MWG Eurofins (Ebersberg, Germany). Genevestigator (www.genevestigator.com) (1) and eFP browser (www.bar.utoronto.ca/efp/cgi-bin/efpWeb.cgi) (2) were used for gene expression analysis of public data. Fluorescent signals were detected using a Leica DM6000B/SP8 confocal laser scanning microscope (Leica Microsystems, Wetzlar, Germany) following the recommended laser settings. Histochemical GUS staining was performed using *Pro_RD26_:GUS* reporter lines as described (3). For hormone- and stress-related studies, two-week-old seedlings grown on half-strength MS medium plates were transferred to liquid MS medium containing relevant treatment chemicals and incubated for 3 h (unless otherwise indicated), preceding the GUS staining. For the identification of tomato orthologs PLAZA 4.0 (https://bioinformatics.psb.ugent.be/plaza) (4) and the Sol Genomics webpage (https://solgenomics.net/) were employed.

### Biological material

*Arabidopsis thaliana* (L.) Heynh., accession Col-0, was used as wild type (5). *RD26* transgenic lines and T-DNA insertion mutants were reported by (3). *RSL4ox* (Hyg^R^) and *pRSL4:RSL4-GFP* (Kan^R^) seeds were kindly provided by L. Dolan (University of Oxford, United Kingdom) and used for crosses with *RD26Ox #21* and #42 plants (BASTA^R^). Progenies from crosses were screened for several generations on plates containing the respective markers. Transgenic tomato *Solanum lycopersicum* (cultivar Moneymaker) lines expressing *SlRD26* from the EST-inducible promoter (*SlRD26-IOE*) were obtained by Agrobacterium-mediated transformation with plasmid *pER10-SlRD26*.

### Constructs

To construct *pEXPA7:RFP*, the *EXPA7* promoter (−1 to −500 bp upstream of the translational start codon) was amplified from Arabidopsis Col-0 genomic DNA and cloned into vector pUBC-RFP-Dest (6). Primer sequences are given in **Supplementary Table 4**. The construct was transformed *via Agrobacterium tumefaciens*-mediated transformation into Arabidopsis Col-0 and *pRD26:RD26-GFP*/*rd26-1* plants using the floral deep method (7). Plasmid *pER10-SlRD26* for the generation of *SlRD26-IOE* tomato plants was constructed by inserting the *Solanum lycopersicum RD26* coding sequence into the pER10 vector, using GATEWAY cloning.

### cDNA synthesis and gene expression analysis

cDNA synthesis and qRT-PCR were performed as reported (8). Primers used for qRT-PCR were designed using QuantPrime (www.quantprime.de) (9). Three technical replicates were performed for each cDNA sample. PCR reactions were conducted using an ABI PRISM 7900HT sequence detection system (Applied Biosystems Applera, Darmstadt, Germany). *ACTIN2* and *SlACTIN* (*Solyc10g080500*) were used as reference genes in Arabidopsis and tomato, respectively, for data normalization as follows: expression data were normalized by subtracting the mean *ACTIN* Ct value from the Ct value of the gene of interest (ΔCt).

### Root hair phenotyping

To study the effect of ABA on root hair formation in WT, 5-day-old Col-0 seedlings, grown in half-strength MS medium plates as above, were transferred to new ABA-free or ABA-containing (15 µM) half-strength MS medium and plates were kept at an angle of 45 degrees to allow the roots growing within the medium. Five days after transfer, the length of the PR and the root hair phenotype within the first 10 mm of the root tip were recorded using a Leica MZ 12.5 stereomicroscope, a digital camera (Leica DFC420), and a differential interference contrast microscope (BX51, Olympus). Root hair length was determined from the digital images using ImageJ (https://imagej.nih.gov). At least 15 seedlings from each genotype were used for phenotyping of root hairs. To determine root hair density, ten serial root hair cells per seedling were scored in the maturation zone using a differential interference contrast microscope (BX51, Olympus). Ten seedlings of each genotype were measured.

Five-day-old *RD26-IOE* and *pER8-EV* seedlings grown on half-strength MS medium plates were transferred to new plates containing either ABA (10 µM) or EST (15 µM), or both (omitted in mock-treated seedlings), and plates were kept in aforementioned conditions for 5 days. Thereafter, root and root hair phenotyping was performed as described.

Tomato seedlings grown on plates were transferred to MS agar plates containing EST (15 µM) or ethanol (0.15% [v/v]; mock treatment) and plates were kept in aforementioned conditions for 4 days. Phenotyping of root hairs was performed within the first 10 mm to the root tip.

For root phenotyping under osmotic stress conditions, 4-day-old ½ MS agar-grown seedlings were transferred to new plates containing an overlay of PEG 8000 on surface (and PEG-free plates as control) as described (10). Plates were kept vertically for four days in aforementioned growth chamber conditions and then analysed. For testing in liquid culture, two-week-old seedlings were transferred to PEG-free or PEG-containing (250 g/l) liquid MS medium and incubated for 5 days on a rotary shaker. Subsequently, photographs were taken and the root phenotype was analysed.

For expression analysis, one-week-old Col-0 seedlings grown on vertical ½ MS agar plates (pH 5.7, 1% sucrose) were used and after 24 h of each treatment (if not indicated otherwise) and RNA was extracted from roots. For shifting to more acidic pH, seedlings were transferred to new pH 4.6 ½ MS agar plates (or pH 5.7 as control). For iron deficiency treatment, seedlings were transferred to iron-sufficient medium (MS agar plates 1% sucrose, 0.1 mM FeEDTA substituted for iron sulfate) or iron-deficient agar medium (MS agar plates 1% sucrose, 300 mM ferrozine, an iron chelator substituted for iron sulfate). For sulphur deficiency treatment, seedlings were transferred to sulphur-deficient medium (20.6 mM NH_4_NO_3_, 18.8 mM KNO_3_, 1.25 mM KPO_4_H_2_, 5 µM KI, 2.99 mM CaCl_2_, 0.1 mM H_3_BO_3_, 1 µM Na_2_MoO_4_, 0.1 µM CoCl_2_, 0.1 mM Na_2_EDTA, 1.5 mM MgCl_2_, 0.1 µM CuCl_2_, 0.1 mM FeCl_2_, 0.1 mM MnCl_2_, 0.03 mM ZnCl2, 1% sucrose) or to MS agar plates as control and after12 h samples were harvested. For phosphate deprivation experiments, seedlings were transferred to phosphate-deficient agar plates (½ MS with 0.65 mM K_2_SO_4_ instead of 1.25 mM KH_2_PO_4_, 1.0% sucrose) or ½ MS medium as control and roots were harvested after 48 h. For PEG treatment seedlings were transferred to PEG-infused plates (medium water potential ∼ −0.7 MPa) or to ½ MS agar plates (medium water potential ∼ −0.25 MPa) and after 48 h roots were harvested. For salt stress treatments, seedlings were transferred to ½ MS agar plates containing 150 mM NaCl or no extra NaCl as control. For osmotic stress treatments, seedlings were transferred to PEG-penetrated plates (250 g/l for the overlay medium) or PEG-free plates. For auxin treatment, seedlings were transferred to ½ MS liquid medium containing 1 µM indole-3-acetic acid (IAA) or ethanol (0.1% [v/v]; mock treatment) and after 1.5 h samples were harvested.

### Gene codes

Arabidopsis gene codes are: *ACTIN2*, *At3g18780*; *EXPA7*, *At1g12560*; *RD26*, *At4g27410*; *RSL1*, *At5g37800*; *RSL4*, *At1g27740.* Additional gene codes (Arabidopsis and tomato) are given in **Supplementary Tables 1, 2** and **4**.

## References

1. B. D. Gruber, R. F. Giehl, S. Friedel, N. von Wiren, Plasticity of the Arabidopsis root system under nutrient deficiencies. Plant Physiol 163, 161–179 (2013).

2. J. E. Salazar-Henao, I. C. Velez-Bermudez, W. Schmidt, The regulation and plasticity of root hair patterning and morphogenesis. Development 143, 1848–1858 (2016).

3. L. Dolan et al., Clonal relationships and cell patterning in the root epidermis of Arabidopsis. Development 120, 2465–2474 (1994).

4. D. W. Kim et al., Functional conservation of a root hair cell-specific cis-element in angiosperms with different root hair distribution patterns. Plant Cell 18, 2958–2970 (2006).

5. S. K. Won et al., Cis-element- and transcriptome-based screening of root hair-specific genes and their functional characterization in Arabidopsis. Plant Physiol 150, 1459–1473 (2009).

6. C. Grierson, E. Nielsen, T. Ketelaarc, J. Schiefelbein, Root hairs. Arabidopsis Book 12, e0172 (2014).

7. Q. Lin et al., GLABRA2 directly suppresses basic helix-loop-helix transcription factor genes with diverse functions in root hair development. Plant Cell 27, 2894–2906 (2015).

8. B. Menand et al., An ancient mechanism controls the development of cells with a rooting function in land plants. Science 316, 1477–1480 (2007).

9. K. Yi, B. Menand, E. Bell, L. Dolan, A basic helix-loop-helix transcription factor controls cell growth and size in root hairs. Nat Genet 42, 264–267 (2010).

10. A. Bruex et al., A gene regulatory network for root epidermis cell differentiation in Arabidopsis. PLoS Genet 8, e1002446 (2012).

11. J. Schiefelbein, L. Huang, X. Zheng, Regulation of epidermal cell fate in Arabidopsis roots: the importance of multiple feedback loops. Front Plant Sci 5, 47 (2014).

12. S. Datta, H. Prescott, L. Dolan, Intensity of a pulse of RSL4 transcription factor synthesis determines Arabidopsis root hair cell size. Nat Plants 1, 15138 (2015).

13. P. Vijayakumar, S. Datta, L. Dolan, ROOT HAIR DEFECTIVE SIX-LIKE4 (RSL4) promotes root hair elongation by transcriptionally regulating the expression of genes required for cell growth. New Phytol 212, 944–953 (2016).

14. H. T. Cho, D. J. Cosgrove, Regulation of root hair initiation and expansin gene expression in Arabidopsis. Plant Cell 14, 3237–3253 (2002).

15. K. Vissenberg, S. C. Fry, J. P. Verbelen, Root hair initiation is coupled to a highly localized increase of xyloglucan endotransglycosylase action in Arabidopsis roots. Plant Physiol 127, 1125–1135 (2001).

16. N. N. Chandrika, K. Sundaravelpandian, S. M. Yu, W. Schmidt, ALFIN-LIKE 6 is involved in root hair elongation during phosphate deficiency in Arabidopsis. New Phytol 198, 709–720 (2013).

17. N. Vartanian, L. Marcotte, J. Giraudat, Drought rhizogenesis in Arabidopsis thaliana (differential responses of hormonal mutants). Plant Physiol 104, 761–767 (1994).

18. Y. Wang et al., Salt-induced plasticity of root hair development is caused by ion disequilibrium in Arabidopsis thaliana. J Plant Res 121, 87–96 (2008).

19. K. Nakashima, H. Takasaki, J. Mizoi, K. Shinozaki, K. Yamaguchi-Shinozaki, NAC transcription factors in plant abiotic stress responses. Biochim Biophys Acta 1819, 97–103 (2012).

20. M. Nuruzzaman, A. M. Sharoni, S. Kikuchi, Roles of NAC transcription factors in the regulation of biotic and abiotic stress responses in plants. Front Microbiol 4, 248 (2013).

21. B. Xu et al., Contribution of NAC transcription factors to plant adaptation to land. Science 343, 1505–1508 (2014).

22. M. Fujita et al., A dehydration-induced NAC protein, RD26, is involved in a novel ABA-dependent stress-signaling pathway. Plant J 39, 863–876 (2004).

23. L. S. Tran et al., Isolation and functional analysis of Arabidopsis stress-inducible NAC transcription factors that bind to a drought-responsive cis-element in the early responsive to dehydration stress 1 promoter. Plant Cell 16, 2481–2498 (2004).

24. H. Ye et al., RD26 mediates crosstalk between drought and brassinosteroid signalling pathways. Nat Commun 8, 14573 (2017).

25. J. R. Dinneny et al., Cell identity mediates the response of Arabidopsis roots to abiotic stress. Science 320, 942–945 (2008).

26. A. S. Iyer-Pascuzzi et al., Cell identity regulators link development and stress responses in the Arabidopsis root. Dev Cell 21, 770–782 (2011).

27. B. O. Bargmann et al., A map of cell type-specific auxin responses. Mol Syst Biol 9, 688 (2013).

28. I. Kamranfar et al., Transcription factor RD26 is a key regulator of metabolic reprogramming during dark-induced senescence. New Phytol 218, 1543–1557 (2018).

29. L. Duan et al., Endodermal ABA signaling promotes lateral root quiescence during salt stress in Arabidopsis seedlings. Plant Cell 25, 324–341 (2013).

30. J. A. Schnall, R. S. Quatrano, Abscisic acid elicits the water-stress response in root hairs of Arabidopsis thaliana. Plant Physiol 100, 216–218 (1992).

31. F. Gosti et al., ABI1 protein phosphatase 2C is a negative regulator of abscisic acid signaling. Plant Cell 11, 1897–1910 (1999).

32. M. K. Jensen et al., The Arabidopsis thaliana NAC transcription factor family: structure-function relationships and determinants of ANAC019 stress signalling. Biochem J 426, 183–196 (2010).

33. W. Li, P. Lan, Re-analysis of RNA-seq transcriptome data reveals new aspects of gene activity in Arabidopsis root hairs. Front Plant Sci 6, 421 (2015).

34. S. Picelli, Single-cell RNA-sequencing: The future of genome biology is now. RNA Biol 14, 637–650 (2017).

35. T. Denyer et al., Spatiotemporal developmental trajectories in the Arabidopsis root revealed using high-throughput single-cell RNA sequencing. Dev Cell 48, 840–852.e845 (2019).

36. K. H. Ryu, L. Huang, H. M. Kang, J. Schiefelbein, Single-cell RNA sequencing resolves molecular relationships among individual plant cells. Plant Physiol 179, 1444–1456 (2019).

37. S. M. Velasquez et al., O-glycosylated cell wall proteins are essential in root hair growth. Science 332, 1401–1403 (2011).

38. C. Lin, H. S. Choi, H. T. Cho, Root hair-specific EXPANSIN A7 is required for root hair elongation in Arabidopsis. Mol Cells 31, 393–397 (2011).

39. N. Baumberger, C. Ringli, B. Keller, The chimeric leucine-rich repeat/extensin cell wall protein LRX1 is required for root hair morphogenesis in Arabidopsis thaliana. Genes Dev 15, 1128–1139 (2001).

40. A. Diet et al., The Arabidopsis root hair cell wall formation mutant lrx1 is suppressed by mutations in the RHM1 gene encoding a UDP-L-rhamnose synthase. Plant Cell 18, 1630–1641 (2006).

41. A. K. Boron et al., Proline-rich protein-like PRPL1 controls elongation of root hairs in Arabidopsis thaliana. J Exp Bot 65, 5485–5495 (2014).

42. S. C. Fry et al., Xyloglucan endotransglycosylase, a new wall-loosening enzyme activity from plants. Biochem J 282 (Pt 3), 821–828 (1992).

43. R. Yokoyama, K. Nishitani, A comprehensive expression analysis of all members of a gene family encoding cell-wall enzymes allowed us to predict cis-regulatory regions involved in cell-wall construction in specific organs of Arabidopsis. Plant Cell Physiol 42, 1025–1033 (2001).

44. A. Maris et al., Differences in enzymic properties of five recombinant xyloglucan endotransglucosylase/hydrolase (XTH) proteins of Arabidopsis thaliana. J Exp Bot 62, 261–271 (2011).

45. K. Vissenberg et al., Differential expression of AtXTH17, AtXTH18, AtXTH19 and AtXTH20 genes in Arabidopsis roots. Physiological roles in specification in cell wall construction. Plant Cell Physiol 46, 192–200 (2005).

46. M. Tognolli, C. Penel, H. Greppin, P. Simon, Analysis and expression of the class III peroxidase large gene family in Arabidopsis thaliana. Gene 288, 129–138 (2002).

47. L. Valerio, M. De Meyer, C. Penel, C. Dunand, Expression analysis of the Arabidopsis peroxidase multigenic family. Phytochemistry 65, 1331–1342 (2004).

48. J. Shigeto, Y. Tsutsumi, Diverse functions and reactions of class III peroxidases. New Phytol 209, 1395–1402 (2016).

49. T. Kwon et al., Transcriptional response of Arabidopsis seedlings during spaceflight reveals peroxidase and cell wall remodeling genes associated with root hair development. Am J Bot 102, 21–35 (2015).

50. N. Tanaka et al., SRPP, a cell wall protein is involved in development and protection of seeds and root hairs in Arabidopsis thaliana. Plant Cell Physiol 58, 760–769 (2017).

51. M. A. Jones, M. J. Raymond, N. Smirnoff, Analysis of the root-hair morphogenesis transcriptome reveals the molecular identity of six genes with roles in root-hair development in Arabidopsis. Plant J 45, 83–100 (2006).

52. L. Rishmawi, H. Sun, K. Schneeberger, M. Hulskamp, A. Schrader, Rapid identification of a natural knockout allele of ARMADILLO REPEAT-CONTAINING KINESIN1 that causes root hair branching by mapping-by-sequencing. Plant Physiol 166, 1280–1287 (2014).

53. S. Santi, W. Schmidt, Dissecting iron deficiency-induced proton extrusion in Arabidopsis roots. New Phytol 183, 1072–1084 (2009).

54. W. Yuan et al., Arabidopsis plasma membrane H+-ATPase genes AHA2 and AHA7 have distinct and overlapping roles in the modulation of root tip H+ efflux in response to low-phosphorus stress. J Exp Bot 68, 1731–1741 (2017).

55. J. W. Schiefelbein, C. Somerville, Genetic control of root hair development in Arabidopsis thaliana. Plant Cell 2, 235–243 (1990).

56. J. Foreman et al., Reactive oxygen species produced by NADPH oxidase regulate plant cell growth. Nature 422, 442–446 (2003).

57. S. Takeda et al., Local positive feedback regulation determines cell shape in root hair cells. Science 319, 1241–1244 (2008).

58. C. S. Grierson, K. Roberts, K. A. Feldmann, L. Dolan, The COW1 locus of arabidopsis acts after RHD2, and in parallel with RHD3 and TIP1, to determine the shape, rate of elongation, and number of root hairs produced from each site of hair formation. Plant Physiol 115, 981–990 (1997).

59. K. Bohme et al., The Arabidopsis COW1 gene encodes a phosphatidylinositol transfer protein essential for root hair tip growth. Plant J 40, 686–698 (2004).

60. M. J. Kim, D. Ruzicka, R. Shin, D. P. Schachtman, The Arabidopsis AP2/ERF transcription factor RAP2.11 modulates plant response to low-potassium conditions. Mol Plant 5, 1042–1057 (2012).

61. T. Wada, T. Tachibana, Y. Shimura, K. Okada, Epidermal cell differentiation in Arabidopsis determined by a Myb homolog, CPC. Science 277, 1113–1116 (1997).

62. T. Ishida et al., Arabidopsis TRANSPARENT TESTA GLABRA2 is directly regulated by R2R3 MYB transcription factors and is involved in regulation of GLABRA2 transcription in epidermal differentiation. Plant Cell 19, 2531–2543 (2007).

63. M. M. Lee, J. Schiefelbein, WEREWOLF, a MYB-related protein in Arabidopsis, is a position-dependent regulator of epidermal cell patterning. Cell 99, 473–483 (1999).

64. S. Schellmann et al., TRIPTYCHON and CAPRICE mediate lateral inhibition during trichome and root hair patterning in Arabidopsis. Embo j 21, 5036–5046 (2002).

65. C. Bernhardt et al., The bHLH genes GLABRA3 (GL3) and ENHANCER OF GLABRA3 (EGL3) specify epidermal cell fate in the Arabidopsis root. Development 130, 6431–6439 (2003).

66. V. Kirik, M. Simon, M. Huelskamp, J. Schiefelbein, The ENHANCER OF TRY AND CPC1 gene acts redundantly with TRIPTYCHON and CAPRICE in trichome and root hair cell patterning in Arabidopsis. Dev Biol 268, 506–513 (2004).

67. Y. Hwang, H. S. Choi, H. M. Cho, H. T. Cho, Tracheophytes Contain Conserved Orthologs of a Basic Helix-Loop-Helix Transcription Factor That Modulate ROOT HAIR SPECIFIC Genes. Plant Cell 29, 39–53 (2017).

68. A. Wu et al., JUNGBRUNNEN1, a reactive oxygen species-responsive NAC transcription factor, regulates longevity in Arabidopsis. Plant Cell 24, 482–506 (2012).

69. G. P. Xue et al., TaNAC69 from the NAC superfamily of transcription factors is up-regulated by abiotic stresses in wheat and recognises two consensus DNA-binding sequences. Functional Plant Biology 33, 43–57 (2006).

70. B. Rymen et al., ABA suppresses root hair growth via the OBP4 transcriptional regulator. Plant Physiol 173, 1750–1762 (2017).

71. K. Kaufmann et al., Chromatin immunoprecipitation (ChIP) of plant transcription factors followed by sequencing (ChIP-SEQ) or hybridization to whole genome arrays (ChIP-CHIP). Nat Protoc 5, 457–472 (2010).

72. K. Swarup et al., The auxin influx carrier LAX3 promotes lateral root emergence. Nat Cell Biol 10, 946–954 (2008).

## References

Bargmann, B. O. et al. A map of cell type-specific auxin responses. Mol Syst Biol 9, 688, doi:10.1038/msb.2013.40 (2013).

De Rybel et al. (2012) A role for the root cap in root branching revealed by the non-auxin probe naxillin. Nat. Chem. Biol. 8, 798–805.

De Smet et al. (2008) Receptor-like kinase ACR4 restricts formative cell divisions in the Arabidopsis root. Science 322, 594–597 (2008).

Dinneny, J. R. et al. Cell identity mediates the response of Arabidopsis roots to abiotic stress. Science 320, 942–945, doi:10.1126/science.1153795 (2008).

Iyer-Pascuzzi, A. S. et al. Cell identity regulators link development and stress responses in the Arabidopsis root. Dev Cell 21, 770–782, doi:10.1016/j.devcel.2011.09.009 (2011).

Kim, D. W. et al. Functional conservation of a root hair cell-specific cis-element in angiosperms with different root hair distribution patterns. Plant Cell 18, 2958–2970, doi:10.1105/tpc.106.045229 (2006).

Ryu, K. H., Huang, L., Kang, H. M. and Schiefelbein, J. (2019) Single-cell RNA sequencing resolves molecular relationships among individual plant cells. Plant Physiol. 179, 1444–1456.

Vanneste et al. (2005) Cell cycle progression in the pericycle is not sufficient for SOLITARY ROOT/IAA14-mediated lateral root initiation in *Arabidopsis thaliana*. Plant Cell 17, 3035– 3050.

Van Bel, M. et al. (2018) PLAZA 4.0: an integrative resource for functional, evolutionary and comparative plant genomics. Nucl. Acids Res. 46, 1190–1196.

Voß et al. (2015) The circadian clock rephases during lateral root organ initiation in *Arabidopsis thaliana*. Nat. Commun. 6, 7641.

## References

1. T. Hruz et al., Genevestigator v3: a reference expression database for the meta-analysis of transcriptomes. Adv Bioinformatics, 420747 (2008).

2. D. Winter et al., An “Electronic Fluorescent Pictograph” browser for exploring and analyzing large-scale biological data sets. PLoS One 2, e718 (2007).

3. I. Kamranfar et al., Transcription factor RD26 is a key regulator of metabolic reprogramming during dark-induced senescence. New Phytol 218, 1543–1557 (2018).

4. M. Van Bel et al., PLAZA 4.0: an integrative resource for functional, evolutionary and comparative plant genomics. Nucleic Acids Res 46, D1190–d1196 (2018).

5. A. Maris et al., Differences in enzymic properties of five recombinant xyloglucan endotransglucosylase/hydrolase (XTH) proteins of *Arabidopsis thaliana*. J Exp Bot 62, 261–271 (2011).

6. C. Grefen et al., A ubiquitin-10 promoter-based vector set for fluorescent protein tagging facilitates temporal stability and native protein distribution in transient and stable expression studies. Plant J 64, 355–365 (2010).

7. S. J. Clough, A. F. Bent, Floral dip: a simplified method for Agrobacterium-mediated transformation of *Arabidopsis thaliana*. Plant J 16, 735–743 (1998).

8. C. Caldana, W. R. Scheible, B. Mueller-Roeber, S. Ruzicic, A quantitative RT-PCR platform for high-throughput expression profiling of 2500 rice transcription factors. Plant Methods 3, 7 (2007).

9. S. Arvidsson, M. Kwasniewski, D. M. Riano-Pachon, B. Mueller-Roeber, QuantPrime--a flexible tool for reliable high-throughput primer design for quantitative PCR. BMC Bioinformatics 9, 465 (2008).

10. P. E. Verslues, E. A. Bray, LWR1 and LWR2 are required for osmoregulation and osmotic adjustment in Arabidopsis. Plant Physiol 136, 2831–2842 (2004).

